# Whole blood transcriptome profiles of trypanotolerant and trypanosusceptible cattle highlight a differential modulation of metabolism and immune response during infection by *Trypanosoma congolense*

**DOI:** 10.1101/2022.06.10.495622

**Authors:** Moana Peylhard, David Berthier, Guiguigbaza-Kossigan Dayo, Isabelle Chantal, Souleymane Sylla, Sabine Nidelet, Emeric Dubois, Guillaume Martin, Guilhem Sempéré, Laurence Flori, Sophie Thévenon

**Author notes:** **Corresponding author:** Sophie Thévenon.

## Abstract

Animal African trypanosomosis, caused by blood protozoan parasites transmitted mainly by tsetse flies, represents a major constraint for millions of cattle in sub-Saharan Africa. Exposed cattle include trypanosusceptible indicine breeds, severely affected by the disease, and West African taurine breeds called trypanotolerant owing to their ability to control parasite development, survive and grow in enzootic areas. Until now the genetic basis of trypanotolerance remains unclear. Here, to improve knowledge of the biological processes involved in trypanotolerance versus trypanosusceptibility, we identified bovine genes differentially expressed in five West African cattle breeds during an experimental infection by *Trypanosoma congolense* and their biological functions. To this end, whole blood genome-wide transcriptome of three trypanotolerant taurine breeds (N’Dama, Lagune and Baoulé), one susceptible zebu (Zebu Fulani) and one African taurine x zebu admixed breed (Borgou) were profiled by RNA sequencing at four time points, one before and three during infection. As expected, infection had a major impact on cattle blood transcriptome regardless of the breed. The functional analysis of differentially expressed genes over time in each breed confirmed an early activation of the innate immune response, followed by an activation of the humoral response and an inhibition of T cell functions at the chronic stage of infection. More importantly, we highlighted overlooked features, such as a strong disturbance in host metabolism and cellular energy production that differentiates trypanotolerant and trypanosusceptible breeds. N’Dama breed showed the earliest regulation of immune response, associated with a strong activation of cellular energy production, also observed in Lagune, and to a lesser extent in Baoulé. Susceptible Zebu Fulani breed differed from other breeds by the strongest modification in lipid metabolism regulation. Overall, this study provides a better understanding of the biological mechanisms at work during infection, especially concerning the interplay between immunity and metabolism that seems differentially regulated depending on the cattle breeds.

## Introduction

Animal African Trypanosomosis (AAT) represents a serious impediment to livestock development in endemic areas of Africa (Alsan, 2015). This vector-borne disease is caused by blood extracellular protozoan parasites from the *Trypanosoma* genus (e.g., *Trypanosoma congolense, T. vivax* and, to a lesser extent, *T. b. brucei*) mainly transmitted by tsetse flies (genus *Glossina*). It affects about 50 million cattle in 38 countries in humid and sub-humid zones of Africa by causing high morbidity and mortality (Uilenberg, 1998), (Swallow, 2000). The Food and Agriculture Organization of the United Nations estimated its annual cost at $4.5 billion (Budd, 1999), (Mattioli et al., 2004). Up to now, no vaccine is available and the main prophylactic and curative measures are based on the reduction of transmission rates through vector control (Bouyer et al., 2013), and the use of trypanocides in livestock (Meyer et al., 2016).

Interestingly, West African taurine (*Bos taurus taurus*) breeds (AFT), such as long-horn (i.e., N’Dama) and short-horn breeds (e.g., Somba, Baoulé and Lagune) that have lived in West Africa for about 4000 years (Payne & Hodges, 1997), (Hanotte et al., 2002), possess the ability to survive and remain productive in tsetse-infested areas by controlling parasitemia and limiting anemia and body weight loss caused by AAT (Murray et al., 1984), (CIPEA, 1979). AFT are thus called trypanotolerant. In contrast, both zebu (*Bos taurus indicus*) cattle, which arrived more recently in Africa (<2000 YBP) (Loftus et al., 1994), (Bradley et al., 1996), (Hanotte et al., 2002), and European taurine, which were recently introduced to increase African cattle productivity (Seck et al., 2010), are susceptible to AAT (Roberts & Gray, 1973), (Amene et al., 1991), (Doko et al., 1997).

The trypanotolerant character is therefore a remarkable example of livestock adaptation to a selective pressure caused by a pathogenic agent. However, the molecular or biological mechanisms underlying this trait have puzzled researchers for dozens of years, although some immunological and genetic studies provided some clues. Indeed, the first immunological studies pointed out a better adaptive immune response and an earlier monocyte lineage activation in N’Dama compared to African Boran zebu (Authie et al., 1993), (Taylor et al., 1996), (Sileghem et al., 1993). Nevertheless, according to (Naessens et al., 2003), the hematopoietic system, which participates in anemia reduction, was not involved in the control of parasitemia. Moreover, several genetic studies on trypanotolerant and trypanosusceptible breeds revealed the polygenic architecture of trypanotolerance (Murray et al., 1990), (Trail et al., 1991), (Van der Waaij et al., 2003), (Hanotte et al., 2003). Further transcriptomic studies (O’Gorman et al., 2006), (O’Gorman et al., 2009) showed that N’Dama and Boran Zebu exhibited a roughly similar response to infection by trypanosomes, but subtle differences in response intensity or timing were observed, such as a higher IL6 and IL10 expression in Boran Zebu, or an enhanced B cell activation in N’Dama. By integrating genomic and transcriptomic data, (Noyes et al., 2011) proposed TICAM1 and ARHGAP15 as two candidate genes for trypanotolerance. However, these findings have not been confirmed by other studies analyzing gene coding sequences in several breeds (Alvarez et al., 2015), (Alvarez et al., 2016). More experiments were performed in mouse models, in which the possibility to knock out candidate genes allows to accurately assess their contribution to tolerance or pathology (Cnops et al., 2015), (Magez et al., 2006), (Magez et al., 2007), (Onyilagha et al., 2015). Nevertheless, mice are not natural hosts of livestock trypanosomes, and important physiological features that differentiate them from cattle (Taylor & Mertens, 1999), (Morrison et al., 2016) prevent to directly transpose results between the two species.

So far, studies, which investigated immunological, genetic and transcriptomic features of cattle trypanotolerance, have mainly focused on two breeds, the trypanotolerant N’Dama cattle (long-horn taurine originating from Fouta-Djallon in Republic of Guinea) and the trypanosusceptible Boran Zebu (an East-African Zebu). Nevertheless, other West-African taurine cattle are classified as trypanotolerant (CIPEA, 1979), (Akol et al., 1986), (Rege, 1999) among which we recently confirmed, under experimental conditions, the trypanotolerant status of two West African short-horn breeds, i.e., Lagune and Baoulé breeds, compared to N’Dama and to the trypanosusceptible Zebu Fulani (Berthier et al., 2015). We also underlined the intermediate status of Borgou breed, an admixed breed between African short-horn taurine and African zebu. Biological samples collected during this experiment that allowed a clear characterization of the trypanotolerant status of these overlooked breeds provide the opportunity to finely investigate breed-specific modulation of gene expression during infection.

In order to increase knowledge on host-parasite interactions in trypanotolerant and trypanosusceptible cattle breeds during trypanosomosis, we performed a gene expression profiling of blood cells of five cattle breeds infected by *T. congolense* using RNA-seq technology, namely the well-characterized trypanotolerant N’Dama breed, two overlooked trypanotolerant breeds, Baoulé and Lagune, the Borgou crossbred breed, and one trypanosusceptible breed, the Zebu Fulani. In addition, we performed an in-depth functional analysis of the differentially expressed genes. More precisely, we looked for i) breed-specific transcriptomic signatures in blood before infection, ii) main genes and biological functions that responded to infection, whatever the breed, iii) breed-specific transcriptomic profiles during infection, and iv) basal and dynamic transcriptomic profiles that could be associated with trypanotolerance.

## Material and Methods

### Animals, experimental infection and sampling

A total of 39 animals from five West African cattle breeds, i.e., three AFT comprising N’Dama (NDA, 8 animals), Lagune (LAG, 7 animals), and Baoulé (BAO, 8 animals), Zebu Fulani (ZFU, 8 animals), and Borgou (BOR, 8 animals), were experimentally infected by intravenous inoculation of 10^5^ trypanosomes of the *T. congolense* savannah IL1180 strain. This experimental infection, conducted at CIRDES (Burkina Faso) according to a protocol approved by the Burkinabe ethical committee (Project no. A002-2013 / CE-CM), was described in details in (Berthier et al., 2015).

Cattle blood samples were collected at the jugular vein using Tempus™ Blood RNA Tubes (Applied Biosystems™, USA), which allowed the immediate blocking of mRNA transcription and degradation, at four time points: before infection (named DPI.0, DPI for days post-infection) and during the infection at 20 days post-infection (DPI.20), corresponding roughly to the increase in parasitemia, 30 DPI (DPI.30), around the peak of parasitemia, and 40 DPI (DPI.40), at the time of the entrance in the chronic phase of the disease. They were stored 24 hours at +4°C before treatment.

### RNA extraction and RNA-seq libraries preparation

RNAs were extracted from blood samples using the Tempus™ Spin RNA Isolation Kit (Applied Biosystems™, USA) according to the manufacturer’s instructions. RNA was finally eluted using 80 μl RNase-free buffer. Total RNA was quantified using a Nanodrop One (Thermo Fisher Scientific, USA) and its quality checked on a Bioanalyzer 2100 using RNA 6000 nano kit (Agilent Technologies, USA). Samples with RNA integrity number >=8.70 were selected.

RNA-seq libraries were constructed from 120 RNA samples obtained from six cattle per breed at four time points at the MGX platform in Montpellier (France) using the TruSeq Stranded mRNA Library Prep Kit (Illumina) following the manufacturer’s instruction. Briefly, poly-A RNAs were purified using oligo-d(T) magnetic beads from 1 μg of total RNA. The poly-A+ RNAs were fragmented and reverse transcribed using random hexamers, Super Script II (Life Technologies) and Actinomycin D. During the second strand generation step, dUTP substituted dTTP in order to prevent the second strand to be used as a matrix during the final PCR amplification. Double stranded cDNAs were adenylated at their 3’ ends before ligation that was performed using Illumina’s indexed adapters. Ligated cDNAs were amplified following 15 PCR cycles and PCR products were purified using AMPure XP Beads (Beckman Coulter Genomics). Libraries were validated using a Bioanalyzer on a DNA1000 chip (Agilent) and quantified using the KAPA Library quantification kit (Roche).

### Sequencing process

Clustering was performed on a cBot and sequencing on a HiSeq2000 (Illumina). After quantification, the libraries were equimolarly pooled by 12, leading to 10 multiplexes of 12 samples each. Assignment to the multiplexes was done by random blocking on the time point and the breed, in order for each multiplex to contain the four time points (3 samples per time point) and the five breeds. Each pool was denatured, diluted and clustered on three lanes using the Cluster Generation kit v3 (Illumina). Sequencing was performed using SBS kit v3 (Illumina) in single read 50 nt mode. Raw sequencing data were saved as FASTQ files. The quality of the data was assessed using FastQC from the Babraham Institute. Potential contaminants were investigated with the FastQ Screen software (the Babraham Institute).

### Reads’ alignment to the bovine and trypanosome genomes

Reads were jointly mapped to both reference genomes of bovine (EnsemblDB *Bos taurus* UMD 3.1 release 79) and parasite (TritrypDB *Trypanosoma congolense* IL3000 release 9) using STAR aligner (STAR 2.4.0j, (Dobin et al., 2013)). To this end, mapping was performed in four steps: i) the STAR index was generated from a unique multi-FASTA file obtained by concatenation of the bovine and trypanosome sequence files, ii) reads from all libraries were aligned against the indexed reference sequences in order to identify intron-exon junctions, iii) a new STAR index was generated from the unique multi-FASTA file obtained by concatenation of the bovine and trypanosome sequence files and information of intron-exon junctions, iv) a final step of read mapping was performed on the new STAR index. A maximum of three mismatches were allowed and multi-mapping to up to 20 different positions was permitted according to the following parameters: --alignIntronMin 50 --alignIntronMax 500000 --outFilterMultimapNmax 20 --outFilterMismatchNmax 4 --outSAMunmapped Within. Information on reads location on both reference genomes was contained in final BAM format files.

The percentages of reads that mapped uniquely to the bovine or to the trypanosome genome and reads that had multiple matches were checked using Picard tools 1.130 (Broad Institute) and Samtools 1.2 (Li et al., 2009). A large majority of reads was uniquely aligned to the bovine genome (85 to 88% and from 73% to 82% for the samples sharing the index with the PhiX control, see S1 Table). The joint mapping approach was validated by the very low number of reads that mapped to both genomes (bovine and trypanosome), comprised between 17 and 124 (from 0.00004% to 0.0003% of input reads). The percentage of reads uniquely aligned to the trypanosome genome varied greatly between samples, from 3**x**10^−6^ to 2.31%, with a mean of 0.21% (S1 Table). Before the infection (DPI.0), few hundred reads were aligned on the parasite genome, corresponding to a maximum ratio of 3.8×10^−5^ reads (reads uniquely aligned to the trypanosome genome/number of uniquely aligned reads) (S1 Fig). The very low rate of sequences assigned to the parasite genome before infection was considered as negligible and as a background noise of sequencing technology and mapping algorithm (O’Rawe et al., 2015). During the infection (DPI.20, DPI.30, and DPI.40), the number of reads assigned to the trypanosome genome increased and was closely related to parasitemia (S1 Fig). Two animals (BO5 and Z4) did not show any increase in the ratio of reads uniquely aligned to the trypanosome genome.

### Transcript quantification and data normalization

Quantification of gene expression was performed for each library using FeatureCounts (Subread package 1.4.6-p4 (Liao et al., 2014)). Bovine gene annotation was downloaded from Ensembl 79 (sequence UMD 3.1). Reads were assigned at the exon level and counts were summarized at the gene level, using default parameters (-t exon -g gene_id), corresponding to unambiguously assigning uniquely aligned single-end reads and reversely stranded reads (-s 2).

Following the advised workflow of the Bioconductor package EdgeR 3.18.3 (Robinson et al., 2010), (R Core Team, 2018), we first removed lowly expressed genes, and kept genes that had more than one count per million in at least two libraries. Out of the 24,616 bovine genes annotated in Ensembl 79, 13,107 genes went through the filter. Normalization of count data was performed using the Trimmed Mean of M-values normalization (Robinson et al., 2010) and dispersion was estimated using a Cox-Reid profile-adjusted likelihood (Chen et al., 2014), using the Bioconductor package EdgeR 3.18.3 under R 3.4.0 environment (Robinson et al., 2010), (R Core Team, 2018).

### Exploration of the global structuration of bovine count data and sample selection

A two-dimensional scatterplot was launched on the normalized count data to assess the global structuration of 120 samples (Ritchie et al., 2015). It was completed by a Principal Component Analysis (PCA) performed on the log of normalized count per million using mixOmics_6.3.0 (Le Cao et al., 2016) (with center=TRUE and scale=FALSE). The first factorial plan separated the samples according to sampling time point, except for BO5 and Z4 whose samples clustered together (S2 Fig). This observation and the very low ratio of sequences assigned to the trypanosome genome (S1 Fig) supported the hypothesis that the infection process did not occur in BO5 and Z4, in accordance with the phenotypic analysis that revealed transient parasitemia and no anemia in these animals (Berthier et al., 2015). BO5 and Z4 were thus discarded from subsequent analyses. In addition, a Baoulé (BA3) that was detected positive in parasitemia before the experimental infection, contrary to the others that were negative based on diagnostic tests (Berthier et al., 2015), was also excluded from further analyses. The final data set used for statistical and functional analyses contained 108 RNA-seq libraries, corresponding to 27 animals (6 NDA, 6 LAG, 5 ZFU, 5 BOR, and 5 BAO) and four time points per animal.

### Differential expression analyses of bovine genes

The differential expression analysis of bovine genes was carried out using the Bioconductor package EdgeR 3.18.3 under R 3.4.0 environment (Robinson et al., 2010), (R Core Team, 2018), which models gene count data according to a negative binomial distribution and moderates the degree of over-dispersion across genes. A generalized linear model (GLM) was fitted for each gene, and tests for determining differential expression were done using a likelihood ratio test (McCarthy et al., 2012). We considered nineteen contrasts to assess differential expressions between two conditions using GLM (Fig 1).

**Figure 1.**
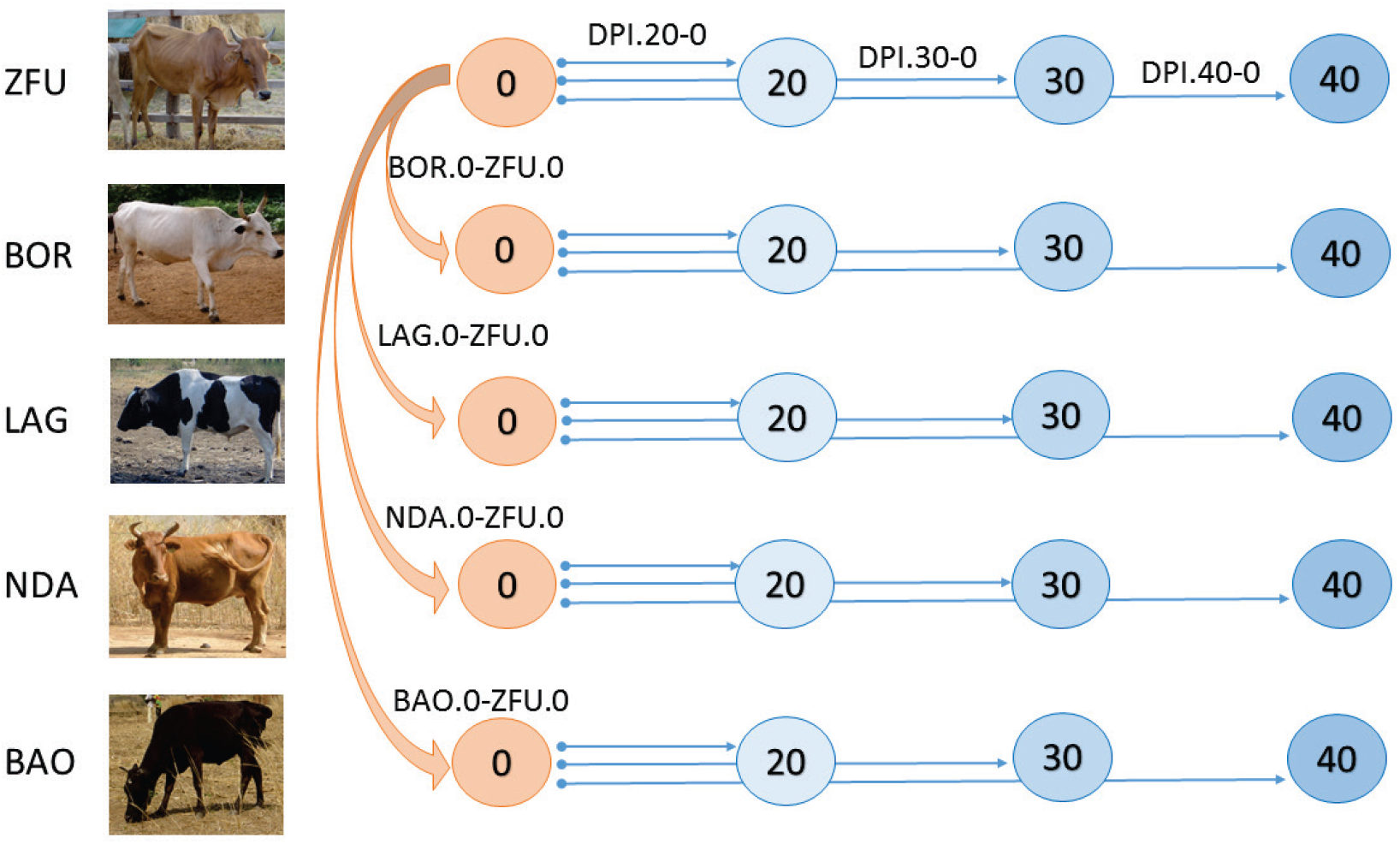
Description of the contrasts used for the differential gene expression analysis. The numbers in the circles represent the days post-infection of cattle sampling, at DPI.0, DPI.20, DPI.30 and DPI.40. Orange arrows represent the contrasts between the breeds at DPI.0, before infection, named BOR.0-ZFU.0, LAG.0-ZFU.0, NDA.0-ZFU.0, and BAO.0-ZFU.0. The blue arrows represent the within-breed contrasts: for each breed, three contrasts were built with DPI.0 as reference, DPI.20-0, DPI.30-0, and DPI.40-0. ZFU, LAG, and BAO pictures by S. Thévenon; BOR picture by G-K. Dayo; NDA picture by D. Berthier.

First, to study baseline differential expression between breeds before infection, we chose ZFU as breed reference, the unique indicine trypanosusceptible breed whose samples were placed at the extremity of the second axis in the PCA (S2 Fig). Four contrasts, comparing NDA, LAG, BAO and BOR breeds to ZFU breed at DPI.0 (Fig 1) were established (namely NDA.0-ZFU.0, LAG.0-ZFU.0, BAO.0-ZFU.0 and BOR.0-ZFU.0). Second, to assess how each breed reacted to infection, 15 within breed contrasts, corresponding to three contrasts that assessed differential expression between three post-infection time points and the pre-infection time point (namely DPI.20-0, DPI.30-0 and DPI.40-0, Fig 1), were considered for each breed. Because animals were repeatedly sampled, the design matrix was constructed according to a nested factorial formula, with animals and time points nested within the breed. GLM likelihood ratio tests provided for each contrast and each gene a logFC (log2-fold change of gene expression between conditions) and a FDR (False Discovery Rate) corresponding to adjusted *p*-values for multiple testing using the Benjamini-Hochberg procedure (Benjamini & Hochberg, 1995). In our analyses, since 19 contrasts were done, a FDR of 10^−3^ (0.05/19=0.0026 rounded to 0.001) was chosen to identify differentially expressed genes (DEGs).

### Functional analysis

The web-based software application Ingenuity^®^ Pathway Analysis (IPA^®^, Version 43605602, 2018-04-04) was used to perform the functional analysis of the DEGs, based on the content of the Ingenuity^®^ Knowledge Base (IKB).

For all contrasts detailed above, among the 13,107 bovine Ensembl identifiers (with their associated logFC and FDR) that defined the background gene list used in IPA^®^ analyses and that were uploaded into the software application, 11,316 identifiers were mapped to their corresponding object in IKB^®^. We then checked via Ensembl Biomart the existence of human orthologues for the 1,791 bovine identifiers that were not recognized in IKB^®^ and found 577 human orthologues with a high confidence level and a one-to-one match. Human Ensembl identifiers for these 577 genes were then used instead of bovine identifiers in files uploaded into IPA^®^. At last, 11,893 Ensembl identifiers were mapped to known genes in IKB^®^.

The functional analysis performed with IPA^®^ identified biological diseases and functions, canonical and signaling pathways and upstream regulatory molecules that were significantly enriched in our data sets. Upstream regulators are regulatory molecules that can affect the expression of target DEGs and that may not have been detected as DEG by RNA-seq (because, e.g., they may be expressed in another tissue than the sampled one, or at another time than the sampling date, or because they are endogenous biochemical compounds). Right-tailed Fisher’s exact test was used to calculate a *p*-value determining the probability that each biological function and/or disease and canonical pathways assigned to our data sets was due to chance alone. These *p*-values were adjusted using the Benjamini-Hochberg correction method (B-H) for diseases and functions and canonical pathways. Diseases and functions, canonical pathways, and upstream regulators were considered as significant if their corresponding *p*-values were below 10^−3^ (B-H correction), 10^−2^ (B-H correction) and 10^−4^ respectively, according to the fact that the quantity of information provided by these functional categories, and thus the number of tests, differed. In addition, based on the logFC of the DEGs and IKB^®^ information, IPA^®^ inferred activation states (namely “activated” or “inhibited”) of biological functions, pathways and upstream regulators (indicating that the observed up or down-regulations of the DEGs are mostly consistent with a particular activation state of a biological function or a regulator) by estimating Z-scores associated with the enriched functions or regulators (Kramer et al., 2014). A Z-score ≥ 2 corresponded to an inferred significant activation state of a function or a regulator, while a Z-score ≤-2 corresponded to an inferred inhibition state. IPA^®^ outputs were visualized using ggplot2 R package (Wickham, 2016).

## Results

In order to better understand host-parasite interactions in trypanotolerant and trypanosusceptible cattle breeds during infection, we carried out an overall assessment of the relationships between the samples, the identification of initially differentially expressed genes between breeds (i.e., before infection), and the identification of differentially expressed genes in each breed during infection, with a first focus on common DEGs and the core associated biological processes, followed by an enlightenment on biological processes specific or prominent to each breed.

### Global overview of bovine gene expression data set

We first carried out an overall assessment of the relationships between the samples, based on a PCA performed on the logarithm of normalized bovine gene counts that provided a global overview of the cattle transcriptomes according to the sampling time point and the breed (Fig 2).

**Figure 2.**
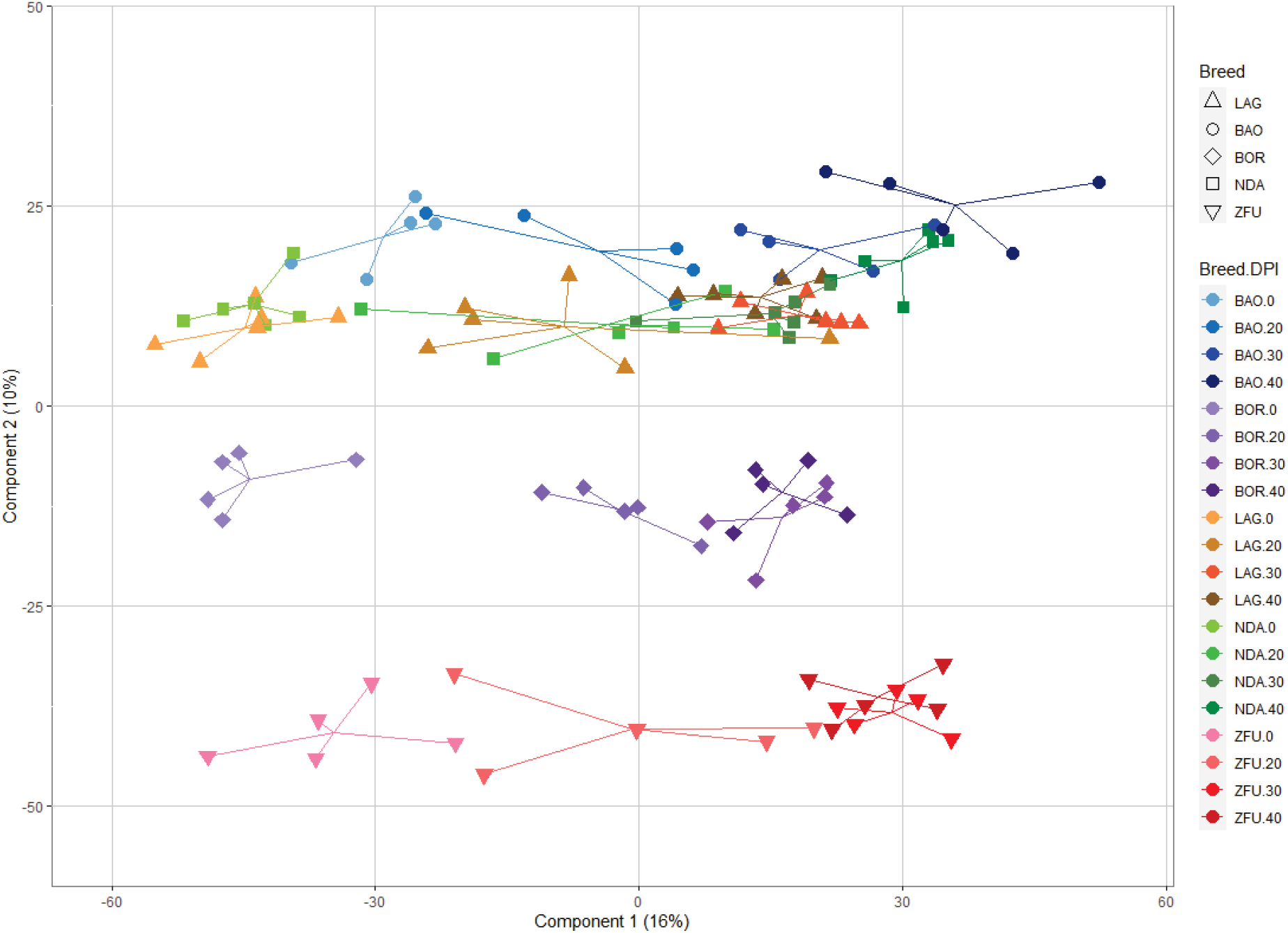
Principal components analysis of cattle RNA-seq libraries based on normalized genes counts. Each point represents a RNA-seq library that corresponds to an animal sampled at a given DPI and that is plotted on the first two principal components according to its coordinates. Libraries are identified according to the breed and the sampling date. Arrows link a library to the centroid of the corresponding breed and sampling date treatment. Each breed is represented by a different shape and by a color gradient, and each color is graded from light to dark shades corresponding to days post-infection respectively DPI.0, 20, 30 and 40. ZFU animals are in red and triangle down, BOR animals are in violet and diamond, LAG animals are in orange and triangle, BAO animals are in blue and circle, and NDA animals are in green and square. Percentages of variance explanation of the first two components are added.

The first axis that accounted for 16% of the total variation was representative of the infection course, from DPI.0 to DPI.40. Shifts from DPI.0 to DPI.40 for each breed were roughly parallel. The second axis that accounted for 10% of the total variation was representative of the breed effect with ZFU count data, at the bottom, taurine count data (represented by NDA, BAO and LAG breeds) at the top and admixed BOR breed data, in the middle, suggesting that basal and lasting count differences existed between zebu, taurine and admixed breeds.

Before infection, a total of 310 genes were detected as differentially expressed (DE) between at least one breed and ZFU (FDR<10^−3^). During infection, a total of 5,270 differentially expressed genes (DEGs) were identified in at least one contrast, between 41 and 3,839 genes being DE depending on the contrast (Fig 3).

**Figure 3.**
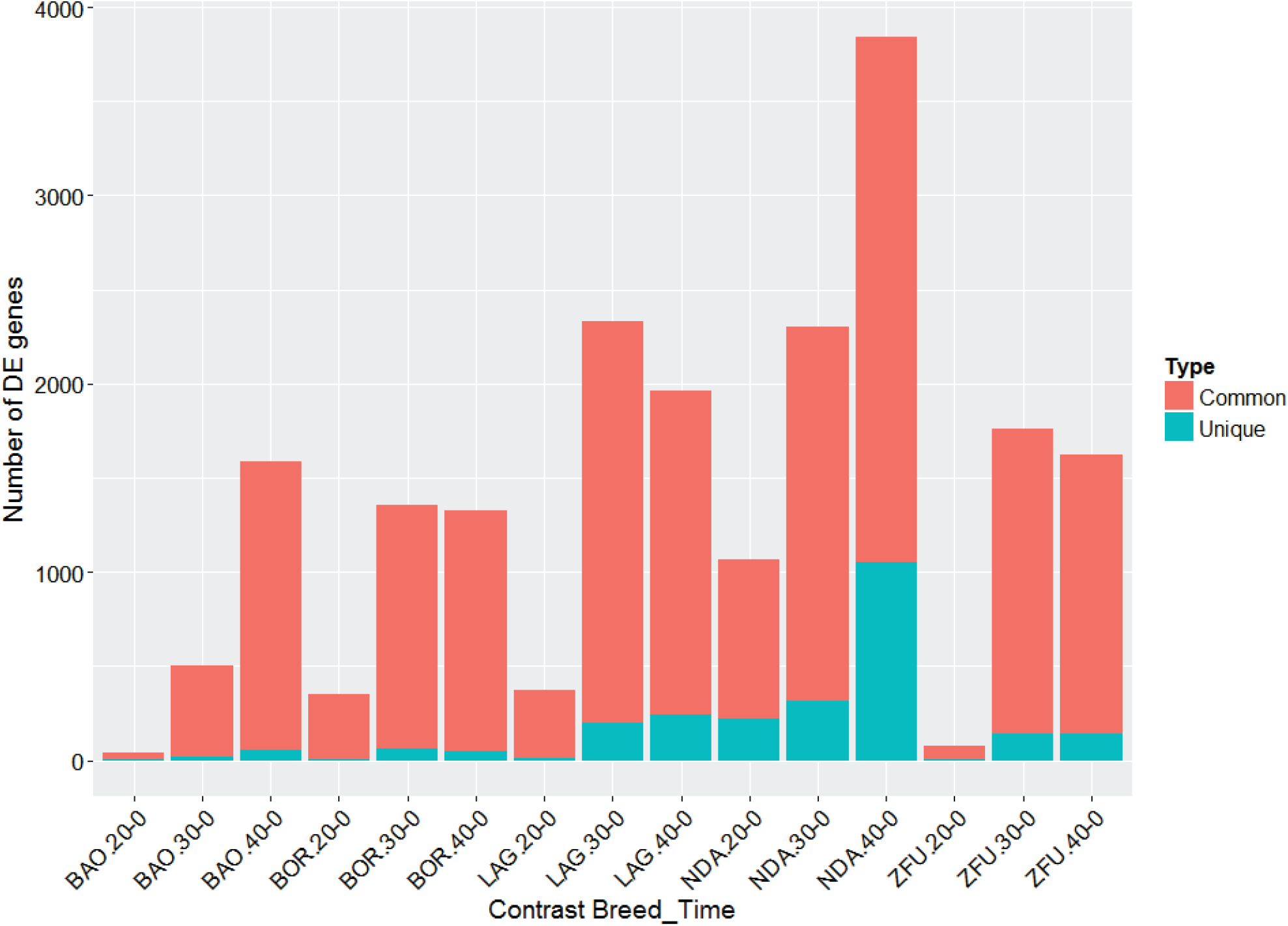
Numbers of genes identified as differentially expressed in the 15 within-breed contrasts. The x-axis represents the 15 contrasts ordered by breed respectively BAO, BOR, LAG, NDA and ZFU, and for each breed, three contrasts according to days post-infection in comparison with before infection, respectively DPI.20-0, 30-0, 40-0. The y-axis represents the number of differentially expressed genes at the threshold of FDR<0.001. The red color corresponds to the part of genes that are differentially expressed and common to, at least, two breeds whatever the sampling time point; the blue color corresponds to the part of genes that are differentially expressed uniquely within a breed.

As expected, the bovine transcriptome was massively modified by trypanosome infection, the total number of DEGs during infection being in descending order 4,344, 2,715, 2,141, 1,753 and 1,696 for NDA, LAG, ZFU, BOR and BAO, respectively. NDA displayed the earliest modulation of its transcriptome, with 1,067 DEGs at DPI.20-0, followed by LAG (370 DEGs), BOR (351 DEGs), ZFU (77 DEGs), and BAO (41 DEGs) (Fig 3). At DPI.30-0, LAG displayed the highest number of DEGs (2,330). At DPI.40-0, NDA showed again the highest number of DEGs (3,839). Many DEGs were common between two or more breeds including the trypanosusceptible ZFU (Fig 3 and S3 Fig). A total of 3,283 genes (62.3% of total number of DEGs, so a majority) were shared between at least two breeds. For instance, 1,823 DEGs were shared between NDA and ZFU, 1,318 between LAG and ZFU, 1,074 between BOR and ZFU, and 1,069 between BAO and ZFU (S3 Fig), whereas the number of DEGs detected exclusively within a breed was 1,272 in NDA, 331 in LAG, 211 in ZFU, 96 in BOR and 77 in BAO. Interestingly, the direction of variation of DEGs, when detected DE in several contrasts, whatever the time point or the breed, was always identical in the aforementioned contrasts (i.e., upregulated with positive logFC or downregulated with negative logFC) except for one gene, SPARC, which was upregulated in LAG.40-0 but downregulated in BAO.40-0 (S2 Table). The heatmap (Fig 4), performed on the logFCs of 5,270 DEGs in at least one contrast, first clustered contrasts according to the time of infection, the early date (DPI.20-0) being separated from the others, except for the contrast BAO.30-0. At later time points, contrasts from a same breed tended to cluster together (LAG.30-0 with LAG.40-0, BOR.30-0 with BOR.40-0, ZFU.30-0 with ZFU.40-0), although NDA.40-0 and BAO.40-0 were not close to NDA.30-0 and BAO.30-0 respectively. Fig 4 did not highlight obvious different patterns of gene expression between breeds.

**Figure 4.**
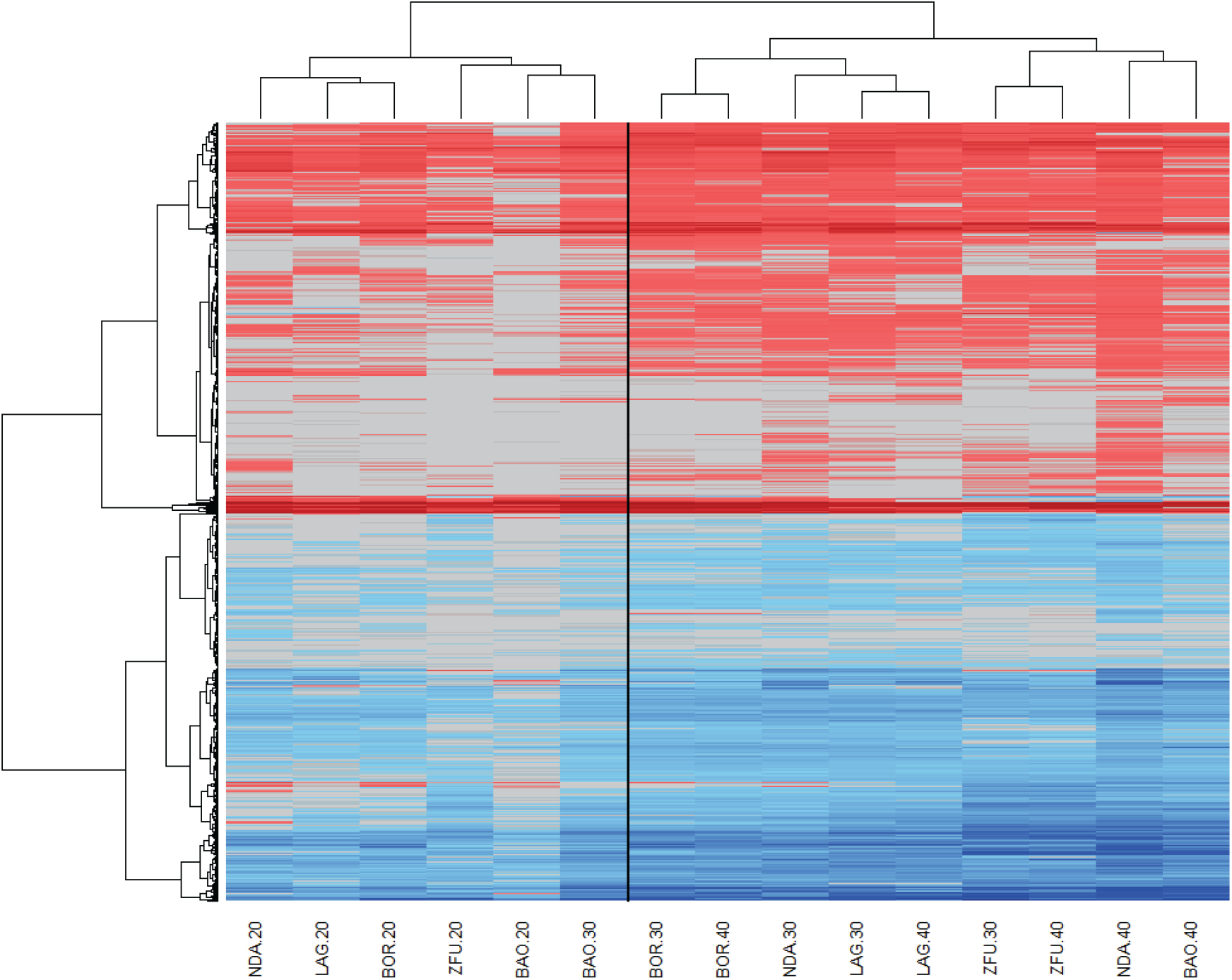
Heatmap on the logFC of 5,270 DEGs in the 15 within-breed contrasts. Columns represent the within-breed contrasts ordered according to clustering, and rows the genes identified as DE in at least one contrast. Up-regulated genes (i.e. with positive logFC) are colored in warm colors from light red to dark red (estimated maximum logFC=11.3), whereas down-regulated genes (i.e. with negative logFC) are colored in cold colors, from light blue to dark blue (estimated minimum logFC=-8.00). LogFC comprised between −0.20 and 0.20 are colored in grey.

### Transcriptome profiling before infection highlights basal differences between breeds

We then focused on the differences before infection of gene expression levels between each breed and ZFU taken as trypanosusceptible reference. For the contrasts LAG.0-ZFU.0, NDA.0-ZFU.0, BAO.0-ZFU.0 and BOR.0-ZFU.0 respectively, 127, 152, 156, and 63 genes were differentially expressed at the FDR threshold of 10^−3^ (Table S3), showing initial differences between gene counts in ZFU and in the other taurine or admixed breeds. As expected from PCA results, the number of DEGs between ZFU and BOR was smaller than those between ZFU and AFT (i.e., NDA, BAO, LAG) at DPI.0. The functional annotation of DEGs for the four contrasts before infection using IPA^®^ indicated that basal differences were not associated with specific biological function enrichment, since neither any disease and function nor any canonical pathway was significantly enriched in these gene sets. However, fifteen upstream regulators, precisely 1, 1, and 15 for the contrasts LAG.0-ZFU.0, BAO.0-ZFU.0 and BOR.0-ZFU.0 respectively, were identified at a *p*-value<10^−4^ (S4 Table). Not any upstream regulator was significantly enriched in NDA.0-ZFU.0. Only BSG, encoding a plasma membrane protein involved in several molecular functions (i.e., cadherin binding, carbohydrate binding, mannose binding, monocarboxylic acid transmembrane transporter activity, and protein binding) was identified as upstream regulator in the three contrasts involving LAG, BAO and BOR. Several upstream regulators linked to immune response (e.g., IL4, TNF, IFNG, Immunoglobulin, CD3 complex, prostaglandin E2, IL10RA) and transcription regulators (e.g., HDAC3, FOXA2, STAT3, ATF3) were enriched in the BOR.0-ZFU.0 gene set.

Among the DEGs before infection, 35, 54, 83 and 18 genes were only DE in LAG.0-ZFU.0, NDA.0-ZFU.0, BAO.0-ZFU.0 and BOR.0-ZFU.0 contrasts, respectively (S4 Fig). Besides, twelve genes were found DE in the four contrasts, 36 were shared between the three AFT breeds versus ZFU.0, and 70 were shared between LAG.0-ZFU.0 and NDA.0-ZFU.0. Among these latter 70 DEGs, all genes except PELI3 were downregulated in LAG and NDA in comparison to ZFU, among which several genes are known to be linked to immune response, especially IL2RA, GBP2, PELI3, DCSTAMP, PTX3, and MARCO (S3 Table).

### Transcriptomic responses common to all breeds, involving immune response and metabolism, are detected during infection

#### Genes differentially expressed during infection in all breeds

To understand the core response of bovine transcriptome to trypanosome infection, we studied the 659 DEGs common to all breeds (namely DEGs in at least one contrast of each breed, S3 Fig). S2 Table gives detailed information on each gene by indicating whether it was DE, in which breed, and its average FDR and logFC. Six genes (NR4A1, CCL22, IFI30, CTSZ, KYNU, IL17REL), involved in immune response, were significantly upregulated in all contrasts, meaning that they were DE within each breed at each time point during infection (S2 Table). Among the top upregulated genes (i.e., harboring the highest average logFC), we found HBM, coding for a hemoglobin subunit (average logFC=6.96, average -log10(FDR)=5.7), ARG1 (average logFC=6.90, average -log10(FDR)=6.0), and several genes involved in immune response like MAPK12, MMP14 and MAPK11. The top downregulated genes (i.e., DEGs displaying the smallest negative logFC) were UNC5A (average logFC =-2.84, average −log10(FDR)=4.7) followed by OVOS2, ELANE, DAB2, and BPI. A quick overview of potentially interesting DEGs associated with immune response allowed highlighting three cytokines (i.e., IL7 upregulated; IL16 and LIF downregulated) and cytokine receptors (e.g., IL1R, IL6R, IL7R, IL20RB, all down-regulated), transcription regulators (e.g., NFKB1 and NFKB2 upregulated), other receptors (e.g., TFRC upregulated), and numerous immune cell antigens, some up-regulated (e.g., CD109, CD180, CD19, CD1A, CD22, CD40, CD72, CD79B, and MME syn.CD10), and others down-regulated (e.g., CD7, CD2, CD226, CD247, CD27, CD3D, CD3E, CD3G, CD40LG, CD99, and ZAP70). Interestingly, several genes known to be involved in metabolism were up-regulated (e.g., HMGCS1, CYP51A1, FDFT1, IDI1, LSS, MVD, SQLE).

#### Enrichment of diseases and functions in the common DEGs

The functional annotation of DEGs common to the breeds using IPA^®^ identified 164 functions and diseases significantly enriched with B-H corrected *p*-values <10^−3^ (S5 Table), and they could be grouped into 24 large categories displayed in Table 1. The major categories were cell-to-cell signaling and interaction, cellular movement, cellular development, cell death and survival, hematological system development and function, cellular function and maintenance, and lipid metabolism. We could also highlight cell-mediated immune response, immunological disease, inflammatory response, and lymphoid tissue structure and development. More precisely, the top ten diseases and functions in terms of B-H corrected *p*-values (<10^−8^) were: proliferation of lymphocytes, proliferation of lymphatic system cells, proliferation of blood cells, proliferation of immune cells, lymphopoiesis, synthesis of sterol, synthesis of cholesterol, quantity of leukocytes, quantity of lymphocytes, and cell movement of lymphatic system cell (S5 Table). IPA^®^ analyses allowed us to assess the activation or inhibition of the enriched diseases and functions based on the Z-score, considered significant if its absolute value was larger or equal to 2. Twenty-four diseases and functions presented significant Z-scores, and surprisingly most had negative Z-scores that could be related to an inhibition state, for instance: lymphocyte homeostasis, adhesion of lymphocytes, migration of lymphatic system cell, and T cell development. Only two diseases and functions had positive Z-score, namely quantity of B-2 lymphocytes and liver lesion.

**Table 1.** Diseases and functions categories enriched in the common DEGs in the within-breed contrasts. The name of diseases and functions categories was indicated, with for each category, the number of significant functions, their average B-H corrected *p*-value, the range of the B-H corrected *p*-values, the average Z-score, and its range. Na: not available.

| Diseases and functions category | Number of functions | Average B-H *p*-value | Range of B-H *p*-values | Average Z-score | Range of Z-scores |
| --- | --- | --- | --- | --- | --- |
| Cell-To-Cell Signaling and Interaction | 28 | 5.7E-05 | $7 \times 10^{-4}$ ; $10^{-8}$ | -1.78 | -2.71 ; 1.13 |
| Cellular Movement | 19 | 1.5E-04 | $9 \times 10^{-4}$ ; $5 \times 10^{-9}$ | -1.45 | -2.66 ; 0.54 |
| Cellular Development | 18 | 1.0E-04 | $7 \times 10^{-4}$ ; $3 \times 10^{-12}$ | -0.34 | -2.08 ; 1.02 |
| Cell Death and Survival | 16 | 1.5E-04 | $9 \times 10^{-4}$ ; $6 \times 10^{-7}$ | -1.09 | -2.36 ; 0.03 |
| Hematological System Development and Function | 15 | 2.5E-04 | $9 \times 10^{-4}$ ; $4 \times 10^{-9}$ | -0.54 | -2.07 ; 2.25 |
| Cellular Function and Maintenance | 9 | 2.4E-04 | $9 \times 10^{-4}$ ; $2 \times 10^{-7}$ | -0.87 | -2.82 ; 0.14 |
| Lipid Metabolism | 8 | 1.1E-05 | $6 \times 10^{-5}$ ; $6 \times 10^{-10}$ | 1.10 | 0.70 ; 1.63 |
| Cancer | 6 | 2.1E-04 | $6 \times 10^{-4}$ ; $5 \times 10^{-6}$ | -0.05 | -0.43 ; 0.33 |
| Cell Morphology | 6 | 1.9E-04 | $97 \times 10^{-4}$ ; $3 \times 10^{-5}$ | -0.73 | -1.00 ; -0.47 |
| Cell-mediated Immune Response | 6 | 1.7E-04 | $5 \times 10^{-4}$ ; $6 \times 10^{-8}$ | -2.11 | -2.65 ; -1.77 |
| Cell Signaling | 5 | 3.5E-04 | $9 \times 10^{-4}$ ; $5 \times 10^{-6}$ | -0.38 | -1.04 ; -0.05 |
| Immunological Disease | 5 | 2.0E-04 | $7 \times 10^{-4}$ ; $3 \times 10^{-6}$ | -1.52 | -2.35 ; -0.67 |
| Inflammatory Response | 5 | 1.5E-04 | $7 \times 10^{-4}$ ; $5 \times 10^{-8}$ | -0.72 | -2.39 ; 1.21 |
| Lymphoid Tissue Structure and Development | 4 | 2.3E-04 | $9 \times 10^{-4}$ ; $5 \times 10^{-8}$ | -0.43 | -1.15 ; 0.11 |
| Connective Tissue Disorders | 2 | 2.2E-05 | $3 \times 10^{-5}$ ; $9 \times 10^{-6}$ | 0.05 | -0.13 ; 0.23 |
| Embryonic Development | 2 | 5.4E-04 | $9 \times 10^{-4}$ ; $2 \times 10^{-4}$ | -0.58 | -1.14 ; -0.02 |
| Inflammatory Disease | 2 | 7.0E-05 | $8 \times 10^{-5}$ ; $5 \times 10^{-5}$ | -0.76 | -2.43 ; 0.90 |
| Organismal Development | 2 | 1.1E-04 | $2 \times 10^{-4}$ ; $2 \times 10^{-5}$ | Na | Na |
| Cellular Growth and Proliferation | 1 | 3.3E-12 |  | -0.21 |  |
| Tissue Morphology | 1 | 2.6E-06 |  | -0.16 |  |
| Molecular Transport | 1 | 3.2E-04 |  | -0.76 |  |
| Gastrointestinal Disease | 1 | 5.3E-04 |  | 2.06 |  |
| Cardiovascular Disease | 1 | 9.6E-04 |  | Na |  |
| Connective Tissue Development and Function | 1 | 9.6E-04 |  | 0.60 |  |

#### Enrichment of canonical Pathways in the common DEGs

Twenty-three canonical pathways were significantly enriched in common DEGs with a B-H corrected *p*-value <10^−2^ (Table 2). This list highlighted several enriched and activated metabolic pathways (Z-score≥2), e.g., the superpathway of cholesterol biosynthesis, the mevalonate pathway, the TCA (TriCarboxylic acid) cycle II, and the superpathway of geranylgeranyldiphosphate biosynthesis I. Mitochondrial dysfunction was also enriched but without inference on the direction of activation. The second class of enriched pathways concerned the immune response including, for instance, role of NFAT in regulation of the immune response, primary immunodeficiency signaling, and regulation of IL-2 expression in activated and anergic T lymphocytes. Two immune response pathways, PI3K signaling in B lymphocytes and B cell receptor signaling, were estimated as significantly activated.

**Table 2.** Canonical and signaling pathways enriched in the common DEGs in the within-breed contrasts. The name of the canonical and signaling pathways is indicated with its B-H corrected *p*-value, its Z-score, and the ratio between the number of DEGs and the number of genes in the pathway. Highlighted cells correspond to Z-scores with an inferred activation state.

| Canonical and signalling pathways | B-H $p$-value | Z-score | Ratio |
| --- | --- | --- | --- |
| Superpathway of Cholesterol Biosynthesis | 5.2E-08 | 3.61 | 0.52 |
| Mevalonate Pathway I | 6.9E-04 | 2.45 | 0.60 |
| TCA Cycle II (Eukaryotic) | 6.9E-04 | 2.83 | 0.40 |
| Superpathway of Geranylgeranyldiphosphate Biosynthesis I (via Mevalonate) | 6.9E-04 | 2.65 | 0.50 |
| Mitochondrial Dysfunction | 1.7E-03 | Na | 0.15 |
| Role of NFAT in Regulation of the Immune Response | 1.7E-03 | 0.22 | 0.15 |
| Primary Immunodeficiency Signaling | 1.7E-03 | Na | 0.32 |
| PI3K Signaling in B Lymphocytes | 1.7E-03 | 2.32 | 0.16 |
| Cholesterol Biosynthesis I | 1.7E-03 | 2.45 | 0.46 |
| Cholesterol Biosynthesis II (via 24,25-dihydrolanosterol) | 1.7E-03 | 2.45 | 0.46 |
| Cholesterol Biosynthesis III (via Desmosterol) | 1.7E-03 | 2.45 | 0.46 |
| Regulation of IL-2 Expression in Activated and Anergic T Lymphocytes | 2.4E-03 | Na | 0.18 |
| Leukocyte Extravasation Signaling | 4.5E-03 | 0.73 | 0.14 |
| Hematopoiesis from Pluripotent Stem Cells | 4.8E-03 | Na | 0.38 |
| T Cell Receptor Signaling | 5.1E-03 | Na | 0.16 |
| Sirtuin Signaling Pathway | 5.2E-03 | -0.78 | 0.12 |
| IL-7 Signaling Pathway | 5.5E-03 | 0.30 | 0.19 |
| iCOS-iCOSL Signaling in T Helper Cells | 5.6E-03 | 0.33 | 0.17 |
| CD28 Signaling in T Helper Cells | 7.1E-03 | -0.30 | 0.15 |
| B Cell Development | 7.1E-03 | Na | 0.33 |
| Systemic Lupus Erythematosus In B Cell Signaling Pathway | 7.1E-03 | -0.21 | 0.12 |
| B Cell Receptor Signaling | 7.1E-03 | 2.68 | 0.13 |
| Phospholipase C Signaling | 8.9E-03 | 0.24 | 0.12 |

#### Enrichment of upstream Regulators in the common DEGs

Seventy-three upstream regulators that can affect the expression of target DEGs were considered as significantly enriched (*P*-value<10^−4^, S6 Table), and the direction of activation was inferred for some of them. The list of upstream regulators referred to chemicals, simple protein or protein complexes. Four endogenous chemicals were identified: cholesterol, which was inferred as inhibited, beta-estradiol, prostaglandin E2 and D-glucose. Nine cytokines were significantly enriched: TNF, IL15, IL4, IL3, IFNG, IL10, CSF1, the latter being estimated as activated, and IL2 and IL7, both estimated as inhibited. Detected protein complexes were also related to immune response (i.e., immunoglobulin, C4BP, TCR, Ige, and NFkB). Twenty-seven transcription regulators were identified, some involved in metabolism (e.g., the activated SREBF2, SREBF1, and PPARGC1B), cell cycle (e.g., TP53 inhibited, SP1) or in pleiotropic functions (e.g., SIRT2 activated). Several transcription factors were linked to immune response (e.g., STAT6 and STAT3, TCF3 inhibited, and BCL6 activated).

### Specific transcriptomic response of each breed during infection

In order to look for potential breed-specific responses during infection, we performed separate enrichment analyses of the lists of DEGs of the 15 within-breed contrasts.

#### Enrichment of diseases and functions in each breed during infection

The analysis identified 642 diseases and functions significantly enriched in at least one of the 15 contrasts (B-H corrected *p*-values <10^−3^) (S7 Table). The most enriched contrast was ZFU.40-0 with 365 significant diseases and functions, while no disease and function were enriched in LAG.20-0. As for the enrichment of the common DEGs, large categories were cellular development and cell cycle, hematological system development and function, immune response, cell signaling, and lipid metabolism. Among these 642 diseases and functions, 135 displayed significant Z-scores for the corresponding contrasts, 17 being considered as activated (Z-score≥2), while a majority (118) was inhibited (Z-score≤-2). We confirmed the results obtained with the analysis of common DEGs, namely a shared inhibition of numerous cellular functions, especially associated with cell-mediated immune response from 30 DPI (e.g., T cell homeostasis). Fig 5 represents both the significance level and the inhibition state of 41 diseases and functions harboring Z-scores smaller than −3 for the 15 contrasts. Few inhibited functions were set up exclusively by one or few breeds such as the differentiation of mononuclear leukocytes, significantly inhibited in NDA.40-0 only. Chemotaxis of myeloid cells was inhibited in BAO.40-0, LAG.40-0, and ZFU.40-0 contrasts but was considered significantly enriched in ZFU only.

**Figure 5.**
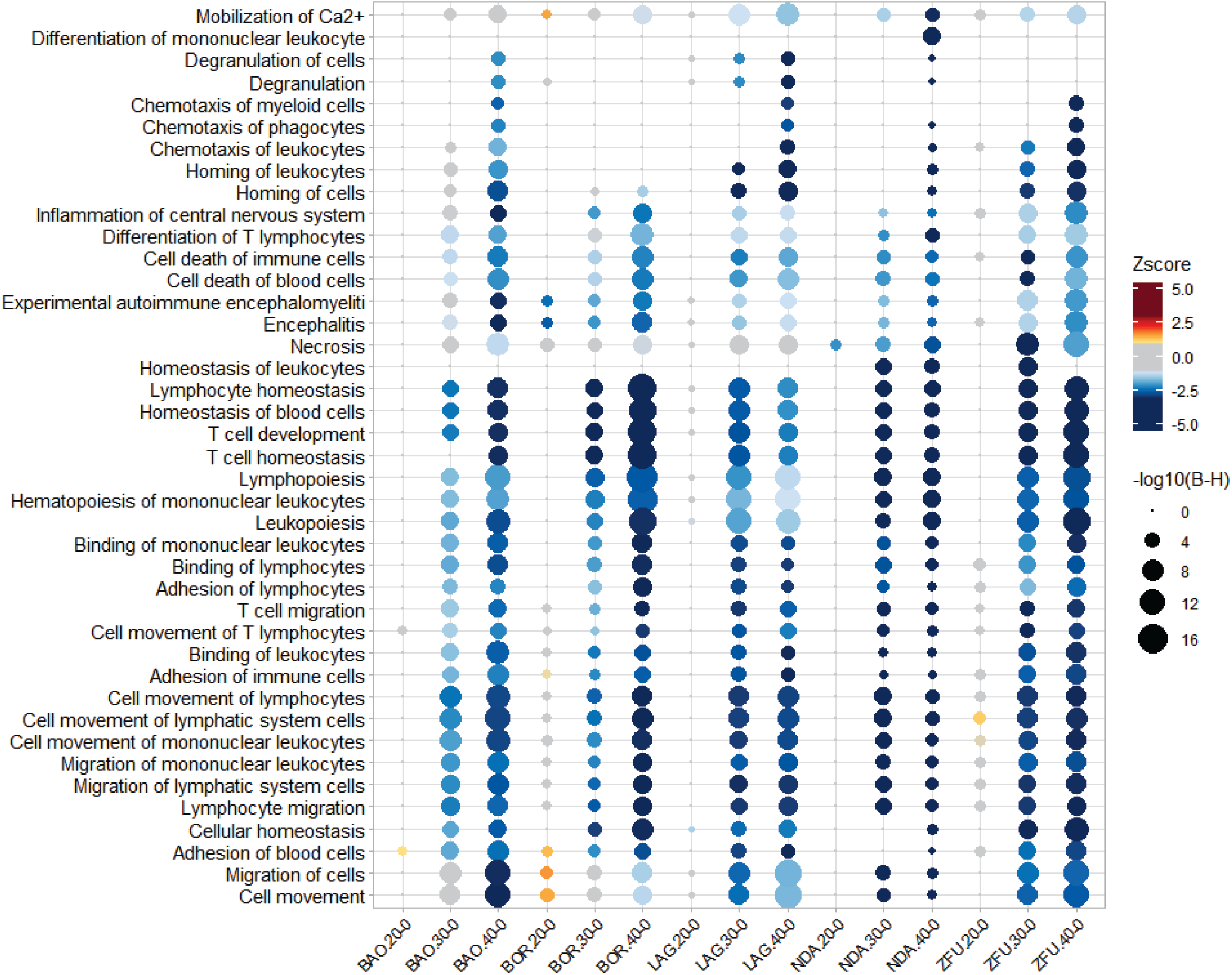
Plot of 41 significantly enriched diseases and functions inferred as inhibited in function of the within-breed contrasts. The 15 within-breeds contrasts are shown on the abscissa and the significantly enriched diseases and functions on the ordinate. The size of the circles is inversely proportional to the B-H corrected *p*-value calculated for diseases and functions in the aforementioned contrast. The color gradient of the circle corresponds to the Z-score range, i.e. warm color gradient for positive Z-score>1, cold color gradient for negative Z-score<-1, and gray color gradient for Z-scores ranging between −1 and 1.

Fewer diseases and functions were activated, but they presented more discriminating patterns than inhibited functions. Fig 6 shows that lipid metabolic functions (synthesis of lipid, synthesis of cholesterol, steroid metabolism, metabolism of cholesterol, and synthesis of terpenoid) were strongly enriched and activated in ZFU.20-0 contrast, to a lesser extent in BAO.30-0, and slightly in NDA.20-0, but were not detected in LAG. Glucose metabolism disorder was enriched in ZFU and BAO, but activated in BAO.30-0 contrast only. Cytopenia was significantly activated in ZFU.40-0 contrast.

**Figure 6.**
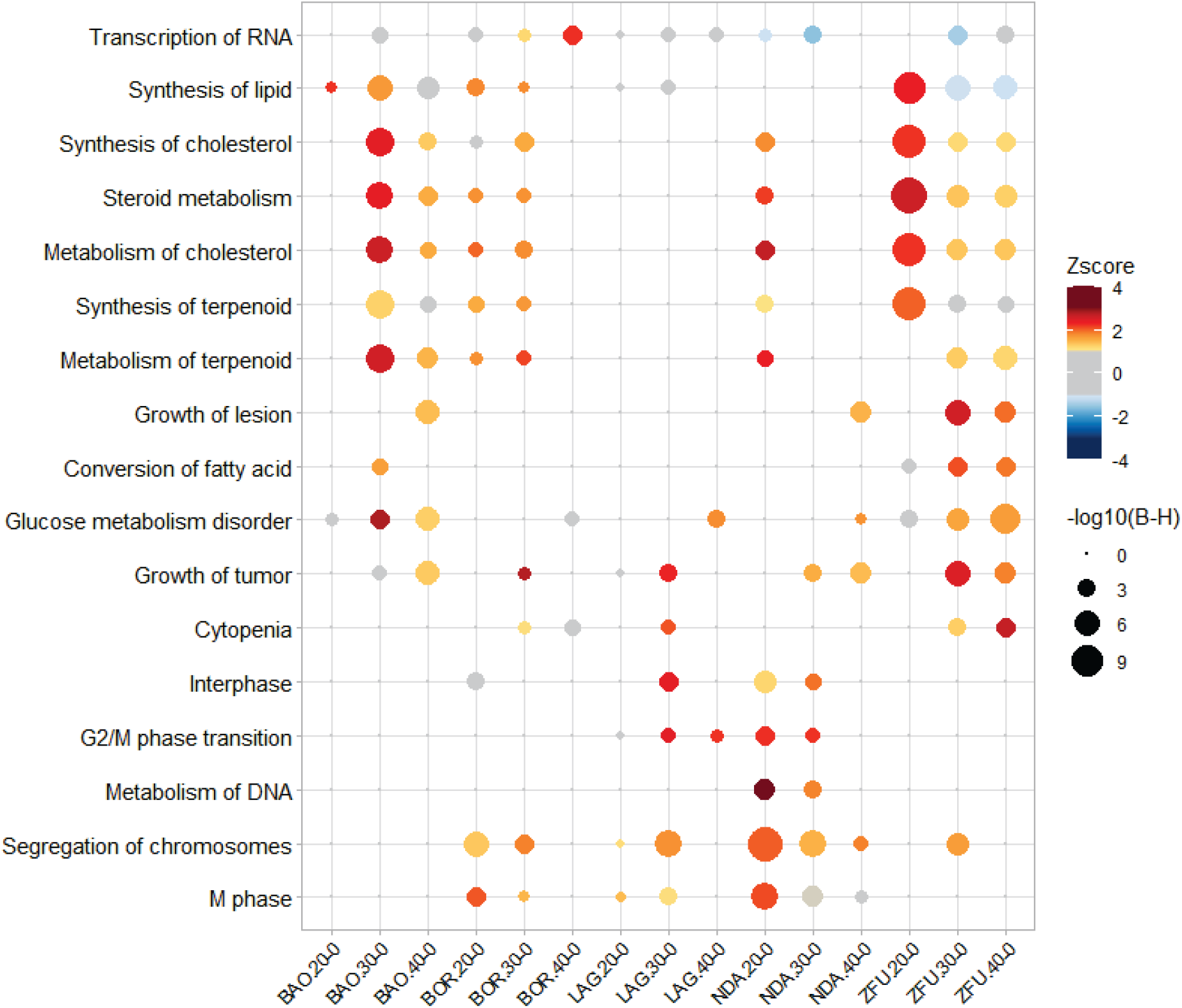
Plot of 17 significantly enriched diseases and functions inferred as activated in function of the within-breed contrasts. The 15 within-breeds contrasts are shown on the abscissa and the significantly enriched diseases and functions on the ordinate. The size of the circles is inversely proportional to the B-H corrected *p*-value of the diseases and functions in the aforementioned contrast. The color gradient of the circle corresponds to the Z-score range, i.e. warm color gradient for positive Z-score>1, cold color gradient for negative Z-score<-1, and gray color gradient for Z-scores ranging between −1 and 1.

The cancer category (i.e., 63 significant functions) and cell cycle functions (i.e., 19 significant functions like cell cycle progression, mitosis) were especially and precociously enriched in NDA.20-0 (S7 Table). G2/M phase transition, metabolism of DNA, segregation of chromosomes, and M phase were strongly activated in NDA.20-0 contrast. This latter was also found activated in BOR.20-0. Interphase was activated in LAG.30-0. BOR.40-0 presented an activation of RNA transcription.

#### Enrichment of canonical pathways in each breed during infection

A total of 92 canonical pathways were significantly enriched in at least one contrast during infection, with a B-H corrected *p*-value<10^−2^ (S8 Table). The contrast ZFU.40-0 displayed the highest number of significant canonical pathways (48), followed by BAO.30-0 (39), while no significant pathways were detected in LAG.20-0, LAG.40-0, and BAO.20-0. Among the 92 significant canonical pathways, 26 presented also significant Z-scores, measuring their activation state: 7 being considered inhibited and 19 activated (Fig 7).

**Figure 7.**
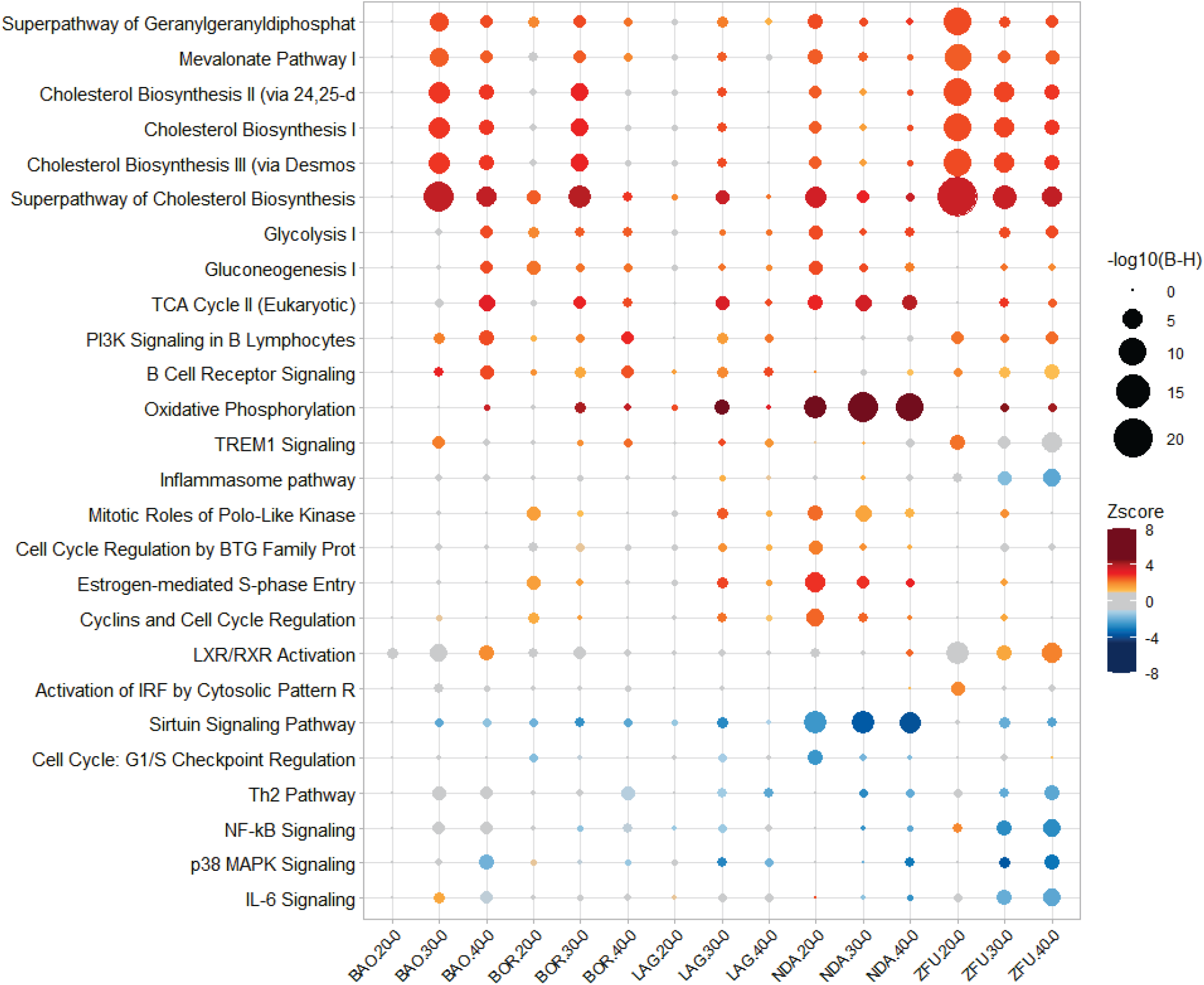
Plot of 26 significantly enriched canonical pathways inferred as activated or inhibited in function of the within-breed contrasts. The 15 within-breeds contrasts are shown on the abscissa and the significantly enriched diseases and functions on the ordinate. The size of the circles is inversely proportional to the B-H corrected *p*-value of the canonical pathways in the aforementioned contrast. The color gradient of the circle corresponds to the Z-score range, i.e. warm color gradient for positive Z-score>1, cold color gradient for negative Z-score<-1, and gray color gradient for Z-scores ranging between −1 and 1.

The superpathway of cholesterol biosynthesis was found significantly activated in all breeds, at one (LAG and NDA), two (BAO and BOR) or three time points (ZFU), but the set of pathways related to cholesterol and lipid metabolisms was particularly and durably enriched in ZFU, and to a lesser extent in BAO. Likewise, LXR/RXR was activated in ZFU.40-0 and almost in BAO.40-0.

Conversely, oxidative phosphorylation, TCA cycle, mitochondrial dysfunction, sirtuin signaling pathway and cell cycle associated pathways were highly significant in NDA contrasts. Pathways associated with cell energy production, oxidative phosphorylation and TCA cycle, were strongly activated in NDA, and to a lesser extent in LAG (oxidative phosphorylation and TCA cycle) and BAO (TCA cycle). Glycolysis and gluconeogenesis were considered enriched and activated in NDA.20-0, and gluconeogenesis was also significantly activated in BOR.20-0. Cell cycle associated pathways (Estrogen-mediated S-phase Entry, Cyclins and Cell Cycle Regulation, Cell Cycle Regulation by BTG Family Proteins, and mitotic Roles of Polo-Like Kinase) were activated in NDA.20-0 exclusively. The sirtuin signaling pathway was highly inhibited in NDA during the infection, and the G1/S checkpoint regulation was inhibited in NDA.20-0.

Some pathways, linked to immune response, were transiently enriched in some breeds, but not in NDA, and their direction of activation was not able to be inferred (i.e., leukocyte extravasation signaling significant in ZFU.40-0, LAG.30-0, and BAO.30-0 and BAO.40-0; hematopoiesis from pluripotent stem cells significant in LAG.30-0, BAO.30-0 and BAO.40-0, and BOR.40-0; natural killer signaling significant in ZFU, BAO and BOR at 40-0). Two B-cell associated pathways (B Cell Receptor signaling, and PI3K Signaling in B Lymphocytes) were activated in BAO.40-0. Interestingly, five pathways were significantly inhibited in ZFU.30-0 and/or ZFU.40-0 (i.e., IL-6 Signaling, NF-kB Signaling, p38 MAPK Signaling, Th2 pathway and the inflammasome pathway).

#### Upstream Regulators enriched in each breed during infection

316 upstream regulators that can affect the expression of target DEGs were considered as significantly enriched in at least one contrast (P-value<10^−4^, S9 Table). Among them, 95 upstream regulators belonged to the transcription regulator category, 29 were endogenous chemicals and 26 were cytokines according to IPA^®^ classification. Upstream regulators found significant in almost all contrasts at the chosen threshold were TGFB1 (14/15 contrasts), TNF (13/15 contrasts), CSF2, Vegf (12/15 contrasts), TCF3, IL4, PTEN (11/15 contrasts). The contrast ZFU.20-0 presented 115 significant upstream regulators, while only eight and 13 were significantly enriched in BAO.20-0 and LAG.20-0 respectively; NDA.20-0 and BOR.20-0 displayed 86 and 87 significant upstream regulators respectively.

In order to visualize the main activated or inhibited regulators predicted by IPA according to the up- or downregulation of the target DEGs, Fig 8 presents the estimated Z-scores of 61 upstream regulators with significant *p*-values (<10^−6^) and Z-scores (|Z-score| ≥2) in at least one significant contrast. Among them, 29 upstream regulators, presenting positive Z-scores, were assessed as activated in some contrasts during infection (top of Fig 8), 25 were roughly inhibited (middle of Fig 8), and 7 showed a dynamic pattern with rather positive Z-scores at DIP.20 followed by negative Z-scores (bottom of Fig 8).

**Figure 8.**
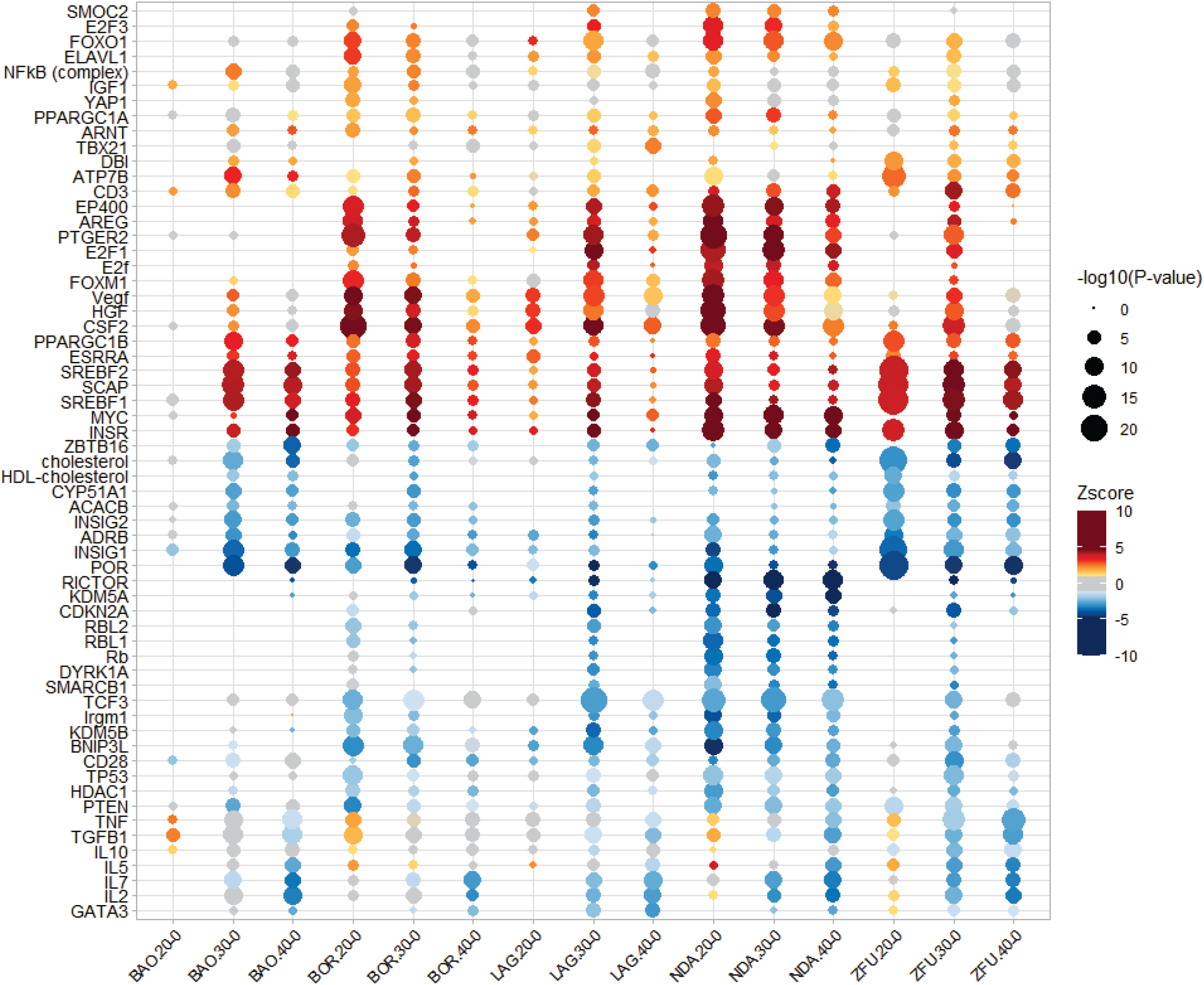
Plot of 61 significantly enriched upstream regulators inferred as activated or inhibited in function of the within-breed contrasts. The 15 within-breeds contrasts are shown on the abscissa and the significantly enriched diseases and functions on the ordinate. The size of the circles is inversely proportional to the P-value of the upstream regulator in the aforementioned contrast. The color gradient of the circle corresponds to the Z-score range, i.e. warm color gradient for positive Z-score>1, cold color gradient for negative Z-score<-1, and gray color gradient for Z-scores ranging between −1 and 1.

ZFU was characterized by very high Z-scores and levels of significance during infection for SREBF2, SCAP, and SREBF1. These upstream regulators were also significantly activated in the other breeds but not constantly during infection. Interestingly, the SREBF2 gene itself was significantly and exclusively up-regulated in ZFU contrasts. ATP7B was durably activated in ZFU, and temporarily in BAO and BOR. ZFU displayed constantly inhibited upstream regulators: cholesterol, CYP51A1, ACACB, INSIG1, INSIG2, and POR.

NDA was distinguished on Fig 8 by strong activation of CSF2, INSR, MYC, HGF, Vegf, and PTGER2. These upstream regulators were also activated in the other breeds but with smaller intensities and durations. The high inhibition of RICTOR was a unique feature of NDA, as well as that of KDM5A. The last three breeds, BAO, BOR, and LAG, did not present striking features. The contrast BAO.30-0 shared similarities with ZFU.20-0, naming high activation of SREBF1, SREBF2, and SCAP, and inhibition of INSIG1 and POR. BAO and BOR presented an activation of NFkB (complex). BOR and LAG displayed similar features to NDA (e.g., strong activation of CSF2 or FOXO1). LAG only showed a significant activation of TBX21 and an inhibition of GATA3 at DPI.40-0.

As upstream regulators, several cytokines (IL2, IL5, IL7, and TNF) and the growth factor TGFB1 showed a dynamic pattern: they presented rather positive Z-scores at DPI.20 and were then assessed as significantly inhibited at 30 or 40 DPI (bottom of Fig 8). TNF was significantly activated in BOR.20-0 and significantly inhibited in NDA.40-0 and ZFU.40-0. Likewise, TGFB1 was activated in BAO.20-0 and inhibited in NDA, ZFU and LAG at DPI.40. IL10 was significantly inhibited in ZFU.30-0.

### Some differentially expressed genes between NDA and ZFU before infection are differentially expressed during infection

Because NDA was the breed with the highest total number of DEGs during infection, and ZFU and NDA presented some distinct enriched biological pathways, we decided to compare DEGs detected in the contrast NDA.0-ZFU.0 to those identified during infection in the corresponding within-breed contrasts. Among DEGs identified in the NDA.0-ZFU.0 contrast, 82 (53%) were also found DE during infection either within NDA or within ZFU. A heatmap performed on the logFC of these 82 genes in the within-breed contrasts shows that the first level of clustering was due to the separation between ZFU and the other breeds, and the second level separated the early infection time point (DPI.20-0) from the others (S5 Fig). The most upregulated genes during infection were MARCO (macrophage receptor with collagenous structure, significantly upregulated in all breeds except ZFU during infection), and ENSBTAG00000022715 (unannotated gene in Ensembl). Conversely, IGF2 (Insulin-like Growth Factor 2) was downregulated in all breeds, but especially in NDA, whereas ANKEF1 was significantly downregulated in NDA only.

Among these genes, 55 were involved in canonical pathways or biological functions that were enriched during infection (S10 Table) and, among them, 25 were present in at least ten functions. Top represented DEGs were IL2RA, which was involved in 276 functions, IGF2 in 142, GATA1 in 95, GPB2 in 52, and MARCO in 58 functions. Thirty-three canonical pathways were concerned, among which oxidative phosphorylation, mitochondrial dysfunction, sirtuin pathway, and Th1 and Th2 pathways. 351 diseases and functions contained DEGs in NDA.0-ZFU.0, top functions in term of number of genes were in particular associated with cancer, cell death and survival, cell movement, cell-to-cell signaling and interaction, hematological disease and inflammatory response. At last, 105 upstream regulators could influence the expression of genes among which some were DE between ZFU and NDA before infection. Top enriched upstream regulators included cytokines (i.e., INFG, IL2, TNF, CSF2, IL4), growth factors (TGFB1, HGF), TP53, Vegf and MYC.

## Discussion

Though bovine trypanotolerance has been described for more than a century (Pierre, 1906), the biological bases of this phenotype remain poorly understood, due to the complexity of the trait that is multigenic and multifactorial (dependent on environmental factors), and to the difficulty to perform experimental infections on cattle, in comparison with model species (Morrison et al., 2016). In order to increase knowledge on the physio-pathology of *T. congolense* infection in cattle, our study provides the first whole blood transcriptome profiling using RNA-seq of five bovine breeds, including African taurine shorthorn and longhorn, during an experimental trypanosome infection. The deep functional analysis of the differentially expressed genes before and during infection identified new clues about trypanotolerance. More precisely, our results confirmed and clarified some previous observations about the genes and the main biological functions modulated during trypanosome infection in cattle (O’Gorman et al., 2006), (O’Gorman et al., 2009). We observed a massive modification of the bovine transcriptome during *T. congolense* infection in five West-African breeds, in accordance with the results of these previous experiments that compared N’Dama and Boran breeds, notwithstanding that the experiments were done on different cells (whole blood in the present study instead of Peripheral Blood Mononuclear Cells in previous studies) using different techniques (RNA-seq in this study versus microarrays and/or qRT-PCR in previous studies). In addition, the six genes (i.e., IL6, CCL2, IL10, IL1RN, IL8, NFKB1) DE in (O’Gorman et al., 2006) and our study varied in the same direction, e.g., IL6 and NFKB1 were clearly upregulated, while IL1RN was strongly downregulated in all breeds (except BOR). Among 23 DE genes representing various biological processes and selected by (O’Gorman et al., 2009) for qRT-PCR validation, sixteen were also DE in the same direction in our data set (eight upregulated and eight downregulated in both experiments, the remaining seven being not significantly expressed in our data set or not annotated in Ensembl).

### A core transcriptomic response in blood cells of all five West-African breeds infected by *T. congolense*

Our global analysis of gene expression of blood cells and of their associated biological functions completed previous works by showing that blood cell transcriptomes responded globally similarly to trypanosome infection whatever the breed. Indeed, all breeds exhibited a parallel shift in their gene expression profiling. A majority of DEGs (62.3%) were shared among at least two breeds, and fold-changes of all DEGs common to two or more breeds varied in the same direction, except one gene (i.e., SPARC). This core cattle transcriptomic response to trypanosome infection is a reminder that cattle, whatever the breed, experience parasitemia waves and show symptoms (e.g., anemia), albeit at varying intensities and durations depending on the breed, as observed in experiments (Berthier et al., 2015), (Authie et al., 1993), (Naessens et al., 2003), (Van der Waaij et al., 2003), or in the field (Trail et al., 1994). The functional annotation of the DEGs shared between breeds during infection also allowed us to provide a global picture of the impact of infection on whole blood gene transcription, by identifying hematopoiesis and immune response, and metabolism as the main gene functions targeted by *T. congolense* infection.

### Modulation of hematopoiesis and immune response during infection

We detected in particular the modulation of gene transcription involved in hematopoiesis during infection such as HBM, a highly upregulated DEG in all breeds. This increase of HBM expression suggests that cattle responded quickly to cope with anemia induced by trypanosomes through an important supply of reticulocytes from the bone marrow into the blood, while very low expression level of hemoglobin subunits is normally found in the blood of healthy cattle (Correia et al., 2018). Expression of TFRC, an important gene for erythropoiesis and also for T and B cell development and proliferation (Ned et al., 2003) was also upregulated.

In addition, many top DEGs detected in all breeds are involved in immune response. Among them, five (i.e., NR4A1, CCL22, IFI30, CTSZ and KYNU) out of the six genes significantly upregulated in all within-breeds contrasts during infection are especially expressed in the monocyte lineage and/or dendritic cells (DC), which initiate the immune response, presenting antigens to T lymphocytes (Wang et al., 2018), (Nowyhed et al., 2015), (Vulcano et al., 2001), (Arunachalam et al., 2000), (Kos et al., 2005), (Harden et al., 2016), (Obermajer et al., 2008). These cell types are known to be activated during trypanosomosis in cattle (Anosa et al., 1999) and mice (Magez et al., 2007), and to play a decisive role in protection or pathology (Bosschaerts et al., 2009). More specifically, NR4A1, a nuclear receptor with an inhibitory role in Th1 and Th17 cell differentiation and in CD8 T cell expansion (Wang et al., 2018), (Nowyhed et al., 2015) could have a protective role during trypanosomosis by modulating inflammation (Morias et al., 2015). As for IL17REL, it was associated with inflammation and inflammatory diseases in human (Franke et al., 2010).

Additional common upregulated genes and enriched biological functions and upstream regulators underlined the activation of innate immune response and macrophages, reported in cattle (Sileghem et al., 1993), (Anosa et al., 1999), (Taiwo & Anosa, 2000), (Naessens, 2006) and mouse (Bosschaerts et al., 2009), (Kuriakose et al., 2019). Indeed, important genes known to respond to pathogen stimuli and to trigger immune response were found upregulated in all breeds during infection, such as MAPK11 and MAPK12 (Risco et al., 2012), NFKB1, up-regulated in bovine PBMC during trypanosomosis (O’Gorman et al., 2006), and NFKB2. Activation of NF-kB complex was shown in endothelial cells and macrophages during *in vitro* interactions with trypanosomes where it promoted a pro-inflammatory response (Ammar et al., 2013), (Leppert et al., 2007), but NF-kBp50 encoded by NFKB1 was also able to stimulate the production of the anti-inflammatory cytokine IL-10 (Bosschaerts et al., 2011). Interestingly, a strongly up-regulated common DEG was ARG1, which codes for arginase-1, a key enzyme of arginine metabolism mainly expressed in macrophages and associated with ornithine production and inhibition of NO production (Mills et al., 2000). ARG1 activity is induced in macrophages of mice infected by *T. brucei* (Gobert et al., 2000) or *T. congolense* (Noel et al., 2002) and its expression level depends on the genetic background of trypanotolerant and susceptible mice (Duleu et al., 2004), (Kierstein et al., 2006). In mice infected by *T. congolense*, ARG1 is involved in immune suppression induced by myeloid-derived suppressor cells that impair proliferation and INF-g production of CD4 T cells (Onyilagha et al., 2018). Moreover, arginase activity has a direct positive impact on trypanosome growth through ornithine production (Fairlamb & Cerami, 1992), (De Muylder et al., 2013). In our data set, ARG1 was found to be highly expressed during infection, similarly to what is found in mice, but with no straightforward differences between breeds. Activation of DC and macrophages may also be highlighted by down-regulated genes, like DAB2 (a Clathrin Adaptor Protein) that acts as an intrinsic negative regulator of immune function and inflammation in those cells (Ahmed et al., 2015), (Hung et al., 2016).

Macrophage activation was also highlighted by the enrichment of upstream regulators like CSF1 (highly enriched and activated in the common DEGs) and CSF2 (highly enriched and activated in within-breed contrasts), which are essential cytokines enhancing macrophage and DC survival and activation (Becher et al., 2016), IFNG, shown to be essential in the early-stage control of trypanosomosis in the mouse (Magez et al., 2020), and TNF, which changed from a rather activated state at DPI.20-0 to an inhibition state at DPI.40-0 recalling the switch from pro-inflammatory macrophages to anti-inflammatory macrophages highlighted in tolerant mouse models (De Baetselier et al., 2001). Looking globally at the results, common DEGs were rather associated with an anti-inflammatory balance (e.g., ARG1), as exemplified by the function inflammation of central nervous system which was inferred as inhibited. Nevertheless, it is speculative to infer pro-inflammatory or anti-inflammatory macrophages from our data set, as macrophage populations are particularly plastic in space and time, and genes associated with both types were regulated (e.g., CCL22, ARG1, IL1R1, PDC1LG2 regulation fitted with anti-inflammatory macrophages, while IL16, NR1D2, DAB2, IRF5, NFKB fitted with pro-inflammatory macrophages according to (Arango Duque & Descoteaux, 2014), (Jablonski et al., 2015), (Murray, 2017)).

The functional annotation of DEGs common to the breeds also highlighted the enrichment of the activation of B cells (with several upregulated common DEGs highly expressed in B cells, like CD19, CD22, CD40, CD72, CD79B, and CD180), which was previously reported in cattle infected by *T. congolense* (Pinder et al., 1988), (Naessens & Williams, 1992), (Taylor et al., 1996), (O’Gorman et al., 2009), in human (Boda et al., 2009), (Lejon et al., 2014) and mouse (Onyilagha et al., 2015). In addition, we noticed an activation of B Cell Receptor Signaling, PI3K Signaling in B Lymphocytes and Quantity of B-2 cells in which common DEGs were involved, and an enrichment of Quantity of B Lymphocytes in all breeds from DPI.30.

Common DEGs and their functions were also related to T-cell-mediated immunity. We noticed a downregulation of genes mainly expressed in T cells, or NK cells, like CD2, CD3D, CD3E, CD3G, CD7, CD27, CD40L, CD225, CD247, and ZAP70, of IL16, a chemoattractant for CD4 T cells (Wilson et al., 2004), and of LIF (Leukemia Inhibitory Factor), a pleiotropic cytokine expressed by T cells, especially CD4 T cells in human and regulatory T cells in mouse (Metcalfe, 2011). T cell functions were inferred as strongly inhibited (e.g., T cell development, activation of T lymphocytes, T cell migration), especially from DPI.30. Moreover, IL2, an upstream regulator of the common DEGs, was significantly inhibited from DPI.30 in NDA, ZFU and LAG, and from DPI.40 in BAO and BOR, as previously observed in PBMC from infected NDA and Boran Zebu (O’Gorman et al., 2006). This global result is in line with an impairment of T cell functions as previously observed in infected cattle ((Flynn & Sileghem, 1991), (Sileghem & Flynn, 1992b)), suggested in HAT (Boda et al., 2009), and reported in mouse (Uzonna et al., 1998).

If we underlined an inhibition of T cell functions during the chronic phase of the disease, our results also suggest an activation in the earlier phase, as illustrated by the detection of an early activation of the T cell co-receptor CD3 as an upstream regulator. Interestingly, CD3 gene expression may be negatively regulated during T cell activation (Paillard et al., 1990), (Badran et al., 2005), (Krishnan et al., 2001) likely by a negative feed-back system necessary to control this process. In addition, another surprising result was the inferred inhibited state of IL7 as an upstream regulator of the common DEGs during infection, though, at the mRNA level, IL7 expression was significantly and sustainably upregulated and IL7RA was significantly downregulated, which is expected under T cell activation (Alves et al., 2008). Two hypotheses may be proposed. First, since trypanosomes are capable of long-term survival in the vertebrate host thanks to antigenic variation (Jackson et al., 2012), (Matthews et al., 2015), a reduced expression of IL7RA and a resulting hyporesponsiveness to IL7 could constitute a protective way to avoid T cell over-activation. Second, IL7 upregulation could be a consequence of T cell lymphopenia (Fry et al., 2001) due to trypanosome factors (Cnops et al., 2015). Whether this dynamic is due to an immunosuppression induced by trypanosomes, or a component of the host-parasite equilibrium (Taylor, 1998) to avoid a deleterious sustained inflammatory response cannot be answered.

### Modulation of gene expression involved in metabolism during infection

Among the main functions and biological processes identified as significantly enriched in DEGs detected in all breeds during infection, functions related to metabolism are clearly highlighted. First, regulation of lipid metabolism was highly impacted during infection, and could suggest a direct effect of trypanosomes or a modulation of the host immune response (Kay et al., 2006), (Fessler, 2015). Indeed, several up-regulated genes in all breeds were involved in cholesterol synthesis (e.g., MVD, IDI1, FDFT1, SQLE, LSS, CYP51A1), and accordingly, lipid metabolism functions (e.g., synthesis of cholesterol and sterol), cholesterol biosynthesis and mevalonate pathways, and key upstream regulators of cholesterol synthesis (SREBF1 and SREBF2, INSR) were strongly enriched and activated in the common DEGs data set, while cholesterol, as an upstream regulator, was considered inhibited, thus promoting its own synthesis. If previous studies suggested the involvement of lipid metabolism regulation in the physiopathology of AAT in mice (Kierstein et al., 2006), in cattle and goat infected by *T. congolense* (Traore-Leroux et al., 1987), (Meade et al., 2009), (Rajavel et al., 2020), (Ndoutamia et al., 2002), this is the first time that this involvement has been detected to such an extent. Modification in lipid metabolism could be due to a direct effect of trypanosomes that are able to internalize lipoproteins (Green et al., 2003), but it could also likely reflect an activation of the inflammatory response as suggested by (Bouvier-Muller et al., 2017).

Second, the change observed in our study in the energy production of the cells probably reflects the modification of immune cell energetic metabolism required to support cell growth, proliferation and effector functions (Donnelly & Finlay, 2015) during infection by *T. congolense*. The tricarboxylic acid (TCA) cycle, which is a hub for generating energy and building blocks for macromolecule synthesis as well as releasing intra-cellular signaling molecules (Martinez-Reyes & Chandel, 2020), was indeed highly enriched and activated in the common DEGs that we detected in our study and in some within-breeds contrasts. Besides, gluconeogenesis and glycolysis were also assessed as activated during infection. These two opposite pathways shared many genes whose corresponding proteins may catalyze reactions in two directions. Nevertheless, since PKLR, involved in a key step for glycolysis, was upregulated in all the breeds during infection, and that FBP1, responsible for final steps for gluconeogenesis (Lebigot et al., 2015), was downregulated, we can suppose that glycolysis was actually the activated pathway during infection. These results are concordant with the absence of gluconeogenesis in blood cells and with the interdependency of glycolysis and TCA cycle, glycolysis supplying TCA cycle with pyruvate and providing molecules for synthetizing nucleotides, glycerol and amino acids (Donnelly & Finlay, 2015).

### Breed-specific whole blood transcriptomic responses

A focus on how each breed reacted to infection allowed observing some variations in their transcriptome, in regard to the enrichment and intensity of activation or inhibition of some biological functions. As previously reported by (O’Gorman et al., 2009), the blood transcriptome of NDA cattle, the trypanotolerant breed of reference (Murray et al., 1984), (Hanotte et al., 2003), seemed to respond earlier and more intensely to infection, with a higher number of detected DEGs in this breed during our experiment in comparison with the other breeds, except at DPI.30-0 when LAG had a little more DEGs. In addition, NDA harbored a more intense and earlier activation of several upstream regulators, notably involved in immune response, in comparison with the other breeds, as exemplified by CSF2, the top activated upstream regulator in NDA at DPI.20-0, that is a key cytokine produced by various cells during an infection or an inflammation and involved in monocyte, macrophage, granulocyte and DC functions (Van de Laar et al., 2012), (Hamilton & Achuthan, 2013). The highly estimated activation state of CSF2 in NDA in our study could reflect an earlier and greater activation of macrophages in this breed, consistent with the earlier release of co-stimulatory cytokines by NDA monocytes in comparison with Boran Zebu monocytes (Sileghem et al., 1993) and the hypothesis of an earlier pro-inflammatory response in PBMC from NDA (O’Gorman et al., 2006). PTGER2, the receptor of prostaglandin E2, an inflammation mediator (Kawahara et al., 2015) associated with a pro-inflammation profile in a mouse model of Chagas disease caused by *T. cruzi* (Guerrero et al., 2015), was also considered as highly and precociously activated. In NDA.30-0 and NDA.40-0 contrasts, MYC, activated in M2 macrophages (Pello et al., 2012), (Jablonski et al., 2015), and critical in T cell proliferation and growth following activation (Wang et al., 2011), was the most activated upstream regulator. We could thus hypothesize that trypanotolerance in NDA could be linked to the precocity and the chronology of activation state of cells, which would allow an efficient immune response while avoiding immune disorders (Vincendeau & Bouteille, 2006).

In addition, biological processes related to cell cycle and DNA metabolism (e.g., diseases and functions like segregation of chromosomes, metabolism of DNA; canonical pathways like cyclins and cell cycle regulation; upstream regulators like E2F1, FOXM1), were strongly enriched and activated in blood cells of NDA breed at DPI.20-0 and DPI.30-0, and explained the enrichment of functions linked to cancer. These results suggest an early activation of division and proliferation of some blood cell types in NDA (e.g., lymphocytes, (Naessens & Williams, 1992), (Naessens et al., 2003)). Curiously, like in all breeds, the lymphopoiesis function was assessed as strongly inhibited from DPI.30-0. These results could seem paradoxical but the sets of genes associated with lymphopoiesis on the one hand, and with cell cycle on the other hand, are different, the former function comprising DEGs encoding T cells surface antigens, cytokines receptors and signal transducers, and the latter comprising DEGs involved in mitosis and centromere formation.

Lastly, NDA displayed a particularly important shift in energetic metabolism of blood cells, whose fine tuning is associated with cell type and fate (O’Neill et al., 2016). Indeed, a key feature of NDA canonical pathways was the strong enrichment of oxidative phosphorylation and mitochondrial dysfunction, the former being highly activated such as TCA cycle and glycolysis. The fact that these three pathways were inferred as activated corroborates the presence of diverse immune cell populations and/or B cell activation. Indeed, glycolysis is rather associated with pro-inflammatory macrophages (Mills et al., 2016), activated DCs or effector T cells (McGettrick & O’Neill, 2013), while oxidative phosphorylation is rather associated with anti-inflammatory macrophages, Tregs or memory T cells (McGettrick & O’Neill, 2013), (Mills et al., 2016), and both seem increased in activated B cells (Caro-Maldonado et al., 2014). However, the strongest activation of oxidative phosphorylation observed in NDA could also be associated with a predominance of an anti-inflammatory component (Pearce & Pearce, 2013) during the chronic phase of infection at 30 and 40 DPI. Regulation of metabolic processes was also highlighted by the enrichment of the sirtuin signaling pathway, which was particularly significantly inhibited in NDA. This pathway comprises several proteins involved in ubiquitous processes and especially in a coupling between metabolic and stress factors and inflammatory response (Loftus & Finlay, 2016). In the same way, the most inhibited upstream regulator in NDA was the RICTOR protein that belongs to mTORC2 complex. The latter regulates cell metabolism and is involved in numerous functions, including development (Guertin et al., 2006), insulin signaling by promoting lipogenesis and glycogen synthesis (Yoon, 2017), and immune cell functions.

ZFU, which had the most pronounced anemia, the most durable parasitemia, and the lowest leukocytosis among the five breeds studied (Berthier et al., 2015), was characterized by a high enrichment and a strong activation/inhibition of biological functions, pathways and upstream regulators linked to lipid metabolism from the beginning of infection (i.e., synthesis of cholesterol, superpathway of cholesterol biosynthesis), which could interact with immune response. This was exemplified by SREBF2, a major transcription factor involved in cholesterol metabolism (Bommer & MacDougald, 2011), strongly activated as an upstream regulator, and also continuously upregulated in ZFU exclusively. In addition, the LXR/RXR activation pathway was found significantly enriched in ZFU but not in NDA nor LAG. Interestingly, (Morrison et al., 2010) identified this pathway as differentially enriched between mice infected by two *T. brucei* strains provoking distinct phenotypes. This pathway, notably expressed in hepatocytes and macrophages, is involved in lipid metabolism and innate immunity in macrophages (Joseph et al., 2004), (Ahsan et al., 2018), with rather an anti-inflammatory balance (Schulman, 2017). In the same direction, the significant inhibition of the inflammasome pathway at DPI.40 in ZFU only tended to show an inhibition of the inflammatory response at the chronic stage of infection in this breed (Zamboni & Lima-Junior, 2015).

LAG, whose trypanotolerance has been demonstrated (Berthier et al., 2015) (i.e., mild anemia and quick recovery), showed a similar whole-blood transcriptome profile to NDA during the experiment (Fig 2). If LAG had less DEGs than NDA, it responded relatively intensely to infection, especially at DPI.30-0, and most information about functional analyses was found at this date. In LAG, like in NDA, top activated canonical pathways were linked to cell energy production, i.e., oxidative phosphorylation and TCA Cycle, mitochondrial dysfunction was enriched, and diseases and functions linked to cell cycle functions (G2/M phase transition, Interphase, Segregation of chromosomes) were activated. Superpathway of cholesterol biosynthesis was also activated, but, unlike the other breeds, not any disease and function linked to cholesterol or lipid metabolism was significant. Top upstream regulators were also shared with NDA, as CSF2, the most enriched and activated upstream regulator at DPI.20-0, and HGF, Vegf, MYC, INSR and TCF3. More specific to LAG response, GATA3 and TBX21, reportedly expressed in an opposite way and respectively associated with Th2 and Th1 cells (Chakir et al., 2003), were significantly detected as upstream regulators in LAG.40-0. GATA3, down regulated in all breeds at the mRNA level, was assessed as inhibited in LAG, and TBX21, upregulated in LAG only, as activated. TBX21 was also associated with NK cell development and effector functions (Deng et al., 2015). In LAG, a pro-inflammatory activation of the Th1 cell subset or NK could be considered, but results from the diseases and functions analysis showed rather an inhibition of T cells. Nevertheless, TBX21 being also expressed in B-cells precursors, contrary to GATA3 (Harashima et al., 2005) and being required in B cells for IFNG dependent switching in IgG2a production (Mohr et al., 2010), other biological mechanisms could be involved.

The BAO breed had an unexpected global transcriptomic response in comparison with its trypanotolerant status, given that this breed did not present a significantly different level of anemia from those of NDA and LAG during infection (Berthier et al., 2015). Indeed, the differential expression between the different times after infection and DPI.0 were smaller in BAO than in other breeds, as shown by the PCA visualization (Fig 2) and the small numbers of DEGs detected in BAO during the infection process. This discrepancy between its transcriptomic response and its trypanotolerant status was reflected at the functional analysis. Indeed, the transcriptomic response in BAO showed similar features to that of ZFU regarding DEGs involved in cholesterol and lipid metabolism, while it resembled NDA and LAG regarding TCA cycle activation, control of Ig quantities and B Cell Receptor Signaling.

At last, BOR, an admixed breed between AFZ and AFT (Flori et al., 2014), displayed an expected global transcriptomic response according to its intermediate phenotype between tolerant and susceptible breeds concerning anemia, parasitemia and leucocyte counts (Berthier et al., 2015). The functional analyses of DEGs did not reveal specific responses in BOR, except the activation in BOR.40-0 of RNA transcription. Significant functions were indeed shared with other breeds, such as the M phase and the segregation of chromosomes at 20 DPI, strongly enriched in NDA, or the canonical pathways linked to cholesterol metabolism at DPI.30-0 more enriched in ZFU.

### Baseline transcriptomic differences between trypanotolerant and susceptible breeds point to genes that could influence the outcome of infection

The cross-referencing of the DEG between breeds before infection and within breeds during infection could provide additional information on gene expression and functions associated with trypanotolerance. Though the PCA supports the hypothesis that a majority of the differences in gene expression before infection results from demographic history (see Fig 2), some differences could be adaptive (Whitehead & Crawford, 2006). Indeed, eighty-two genes were DE between NDA and ZFU before infection and also responded to infection in NDA and/or ZFU. Many of these genes are reported to be involved in functions highlighted previously, e.g., immune response and metabolism. One hypothesis is that differences in basal expression of these genes could underline different proportions of cell types or different states of cell activation between breeds, which could play a role in subsequent pathogenic processes caused by trypanosomes.

As illustration, several genes are known to be associated with macrophage functions. Among them, MARCO, whose expression level was higher in ZFU than in NDA and LAG before infection, but was subsequently significantly upregulated in all breeds except in ZFU, is a pattern recognition receptor on macrophages surface involved in phagocytosis of various pathogens and the subsequent enhancement of immune response and chemokines expression (Arredouani et al., 2004), (Bowdish et al., 2009), (Xu et al., 2017). MMD (Monocyte to macrophage differentiation-associated) is a gene expressed during monocyte differentiation and macrophage activation (Liu et al., 2012), and it was similarly upregulated in ZFU.0 in comparison to NDA.0, and upregulated during the infection in NDA, LAG and BOR. SLC11A1 (syn. Nramp1) is also a macrophage gene, and was upregulated in ZFU.0 versus NDA.0, but it was then downregulated in ZFU during infection. The corresponding protein regulates iron homeostasis in macrophages and gene variants are associated with disease susceptibility or resistance (Archer et al., 2015). In addition, IL2RA, which is strongly expressed by Tregs, effector T cells (Banham et al., 2006), NK cells (Esin et al., 2013), (Hamilton et al., 2017), and granulocytes in cattle (Zoldan et al., 2014) was more expressed in ZFU.0 in comparison with the other breeds, and downregulated during infection and particularly in ZFU. IL2RA (syn. CD25) has been the target of research in bovine trypanosomosis, where its impairment was noticed in lymph nodes (Sileghem & Flynn, 1992a). In mice, injection of anti-CD25 antibodies before experimental infections with *T. congolense* led to discordant results related to protection or pathogeny (Okwor et al., 2012), (Guilliams et al., 2007). Other genes harboring basal differences in their expression level were associated with metabolism regulation. For instance, IGF2, which was upregulated in NDA.0 versus ZFU.0 and was downregulated during the infection, codes for a peptide hormone that is involved in metabolism, tissue development and maintenance and is downregulated during under-nutrition or hypoglycemia (Livingstone & Borai, 2014).

In summary, our study provides the first transcriptome profiling of whole blood cells of five West-African bovine breeds during a trypanosome infection using RNA-seq. In order to identify potential similarities among African taurine cattle, overlooked trypanotolerant breeds (i.e., Lagune and Baoule) were considered in addition to N’Dama, which accounts for the majority of the studies about cattle trypanotolerance. We observed that trypanosome infection due to *T. congolense* has a major impact on cattle blood transcriptome, whatever the breed and we provided a global transcriptomic picture of infection. In accordance with previous results, a strong regulation of the immune system functions with an early activation of innate immune response, followed by an activation of humoral response and an inhibition of T cell functions at the chronic stage of infection were detected in all breeds.

Most importantly, we highlighted overlooked features, as a strong modification in lipid metabolism. We mainly noticed an early regulation of the immune response in NDA, associated with a strong activation of energy production by the cell, a strong enrichment and activation of oxidative phosphorylation in NDA and LAG, and an activation of the TCA cycle in AFT breeds, which was not highlighted in ZFU. These differences in cellular energy could be linked in AFT to better functions of some cell types, like M2 macrophages, memory T cells or activated B cells, and could represent a key to decipher trypanotolerance. If some DEGs, functions and biological pathways were shared between AFT breeds during infection, our results highlight also differences in gene expression dynamics in these three trypanotolerant breeds (as exemplified by the singular transcriptomic profile of Baoule). This suggests that AFT breeds, although subjected to the same selective pressure caused by trypanosomes, may have developed different adaptation mechanisms. In addition, the trypanosusceptible breed ZFU presented several canonical pathways linked to inflammation inhibited from DPI.30, which was not observed in the other breeds, and the strongest modification in lipid metabolism regulation. It would be worth exploring other African zebu breeds to confirm if this observation is a global feature of indicine breeds. Some genes, known to be involved in immune response or metabolism, were differently expressed between breeds before infection and within breeds during infection and raise the hypothesis that basal differences between breeds could impact the outcome in infection.

Finally, our study provided new and valuable data to contribute to a better knowledge of African livestock genomics (Kemp, 2019), (Kim et al., 2020), and to decipher the pathogenic process in bovine trypanosomosis due to *T. congolense*. A comparable experiment using *T vivax* or *T. brucei brucei* would be worthwhile to verify whether host responses are similar regardless of the infecting species or not. Overall, interactions between immune response and metabolism deserve to be deeply explored in cattle in order to improve preventive and curative measures of AAT and also other infectious diseases.

## Acknowledgements

This work was supported by the CIRAD - UMR AGAP HPC Data Center of the South Green Bioinformatics platform (http://www.southgreen.fr/). We thank Jean Nakhle for having corrected the English version of the manuscript.

## Data, script and code availability

Raw sequences data (fastq, count matrix, experimental design) will be publicly available at GEO (https://www.ncbi.nlm.nih.gov/geo/) under accession number GSE197108 upon publication.

Scripts and intermediate tables (DEG analysis, input tables for IPA^®^ analyses, output tables from IPA^®^ analyses) are publicly available in Cirad Dataverse under https://doi.org/10.18167/DVN1/L9SHAX. SNP genotypes of the experimental cattle are publicly available in Cirad Dataverse under https://doi.org/10.18167/DVN1/APTZOC and in WIDDE (http://widde.toulouse.inra.fr/widde/).

## Supporting information availability

Tables and figures are in the supplementary tables and figures files. Legends are displayed at the bottom of each tables and figures.

**S1 Table.** Summary information on sequencing and mapping results.

**S2 Table.** LogFC and FDR of the 13,107 genes for the 15 within-breed contrasts during infection.

**S3 Table.** LogFC and FDR of the 13,107 genes for the 4 between-breed contrasts at DPI.0.

**S4 Table.** Upstream regulators significantly enriched in the between-breed contrasts at DPI.0

**S5 Table.** Diseases and functions enriched in the common DEGs in the within-breed contrasts.

**S6 Table.** Upstream regulators enriched in the common DEGs in the within-breed contrasts.

**S7 Table.** Cross table of the enriched diseases and functions in the 15 within-breed contrasts.

**S8 Table.** Cross table of the canonical pathways in the 15 within-breed contrasts.

**S9 Table**. Cross table of the upstream regulators in the 15 within-breed contrasts.

**S10 Table.** Cross table between the functional categories enriched during infection and the genes differentially expressed in the contrast NDA.0-ZFU.0.

**S1 Figure.** Proportion of uniquely mapped reads on the trypanosome genome depending on time points.

**S2 Figure.** Principal components analysis of 120 cattle RNA-seq libraries based on normalized genes counts.

**S3 Figure.** Venn diagram showing the intersection of genes identified as DE in the within-breed contrasts.

**S4 Figure.** Venn diagram showing the intersection of genes identified as DE in the between-breed contrasts.

**S5 Figure.** Heatmap on the logFC of a subset of 82 DE genes.

