## Supplemental Figures for "Whole blood transcriptome profiles of trypanotolerant and trypanosusceptible cattle highlight a differential modulation of metabolism and immune response during infection by *Trypanosoma congolense*"

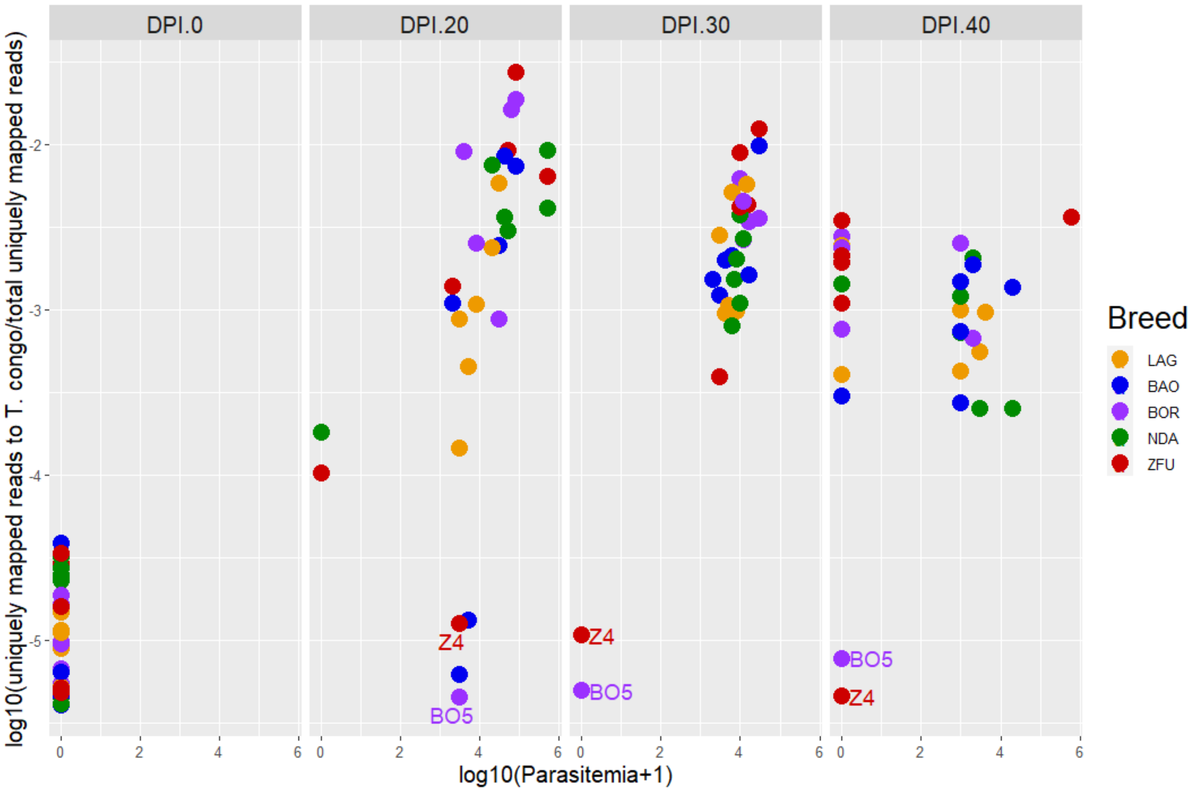


**S1 Figure. Proportion of uniquely mapped reads to the trypanosome genome depending on time points.**

The four panels (DPI.0, DPI.20, DPI.30 and DPI.40) corresponds to the four sampling time points. The x-axis represents the log10 of parasitemia (+1), according to Berthier *et al*., 2015. The y-axis represents the log10 of the proportion of uniquely mapped reads to the trypanosome genome per the total number of uniquely mapped reads (i.e., to trypanosome and bovine genomes). Each point represents a sample, and the colour is function of the breed (yellow for LAG, blue for BAO, violet for BOR, green for NDA and red for ZFU).

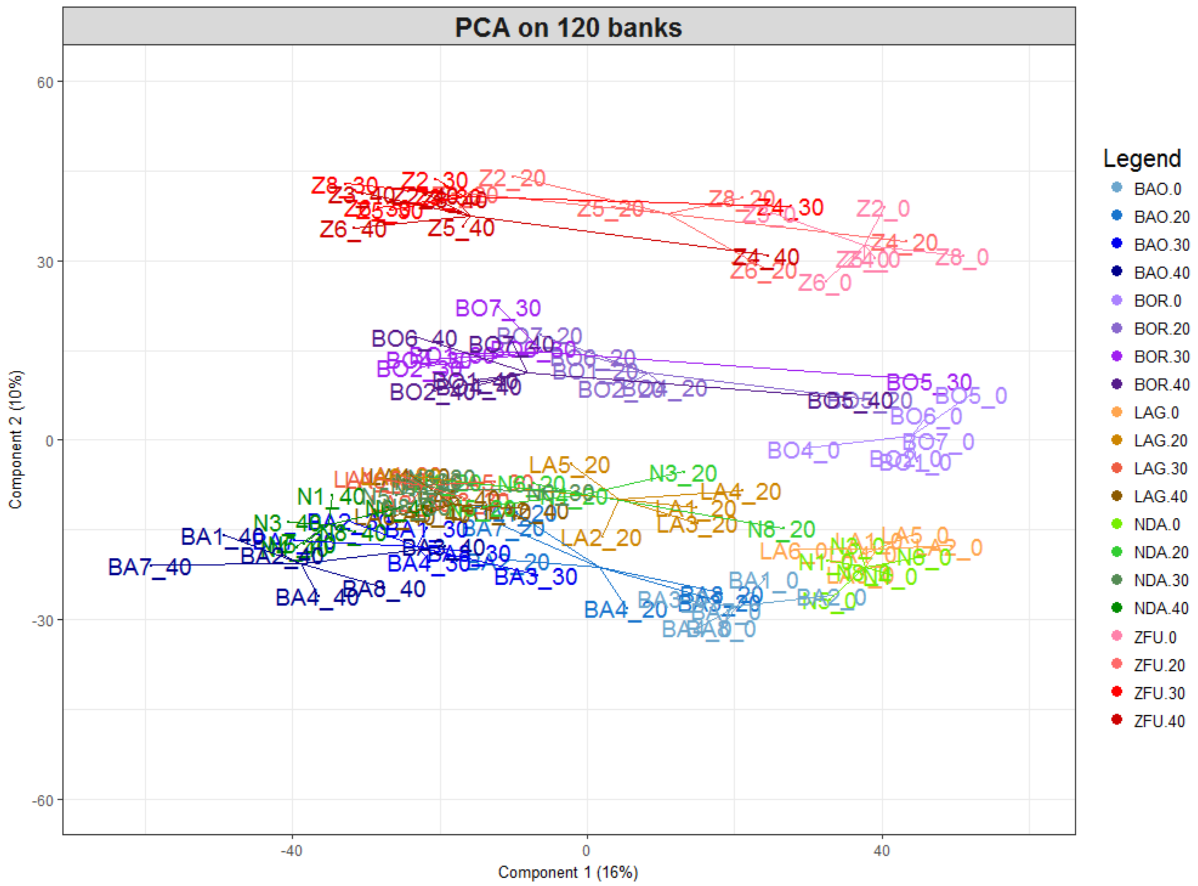


**S2 Figure. Principal components analysis of 120 cattle RNA-seq libraries based on normalized genes counts.**

Each point represents a RNA-seq library that corresponds to an animal sampled at a given DPI and that is plotted on the first two principal components according to its coordinates. Libraries are identified according to the animal identifier and the sampling time point. Each breed is represented by a colour gradient, and each colour is graded from light to dark shades corresponding to days post-infection respectively DPI.0, 20, 30 and 40. ZFU animals (named Z) are in red, BOR animals (BO) in violet, LAG animals (LA) in orange, BAO animals (BA) in blue, and NDA animals (N) in green. Percentages of variance explanation of the first two components are added.

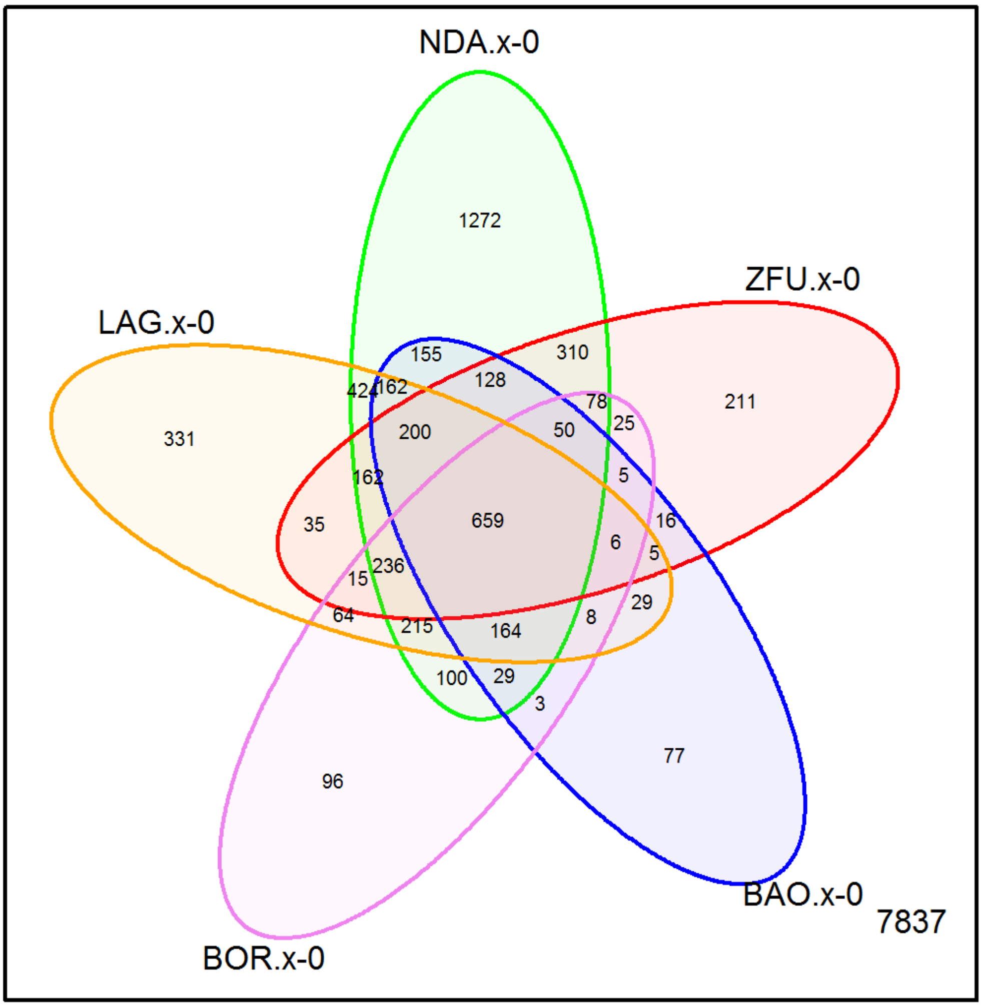


**S3 Figure. Venn diagram showing the intersection of genes identified as DE in the within-breed contrasts.**

Number of genes identified as DE within the five breeds during infection and at the intersections of the different sets. For each breed, the summary of the three contrasts are noticed (i.e., LAG.x-0 represents the number of genes identified as DE in at least one of the three contrasts LAG.20-0, LAG.30-0, and LAG.40-0).

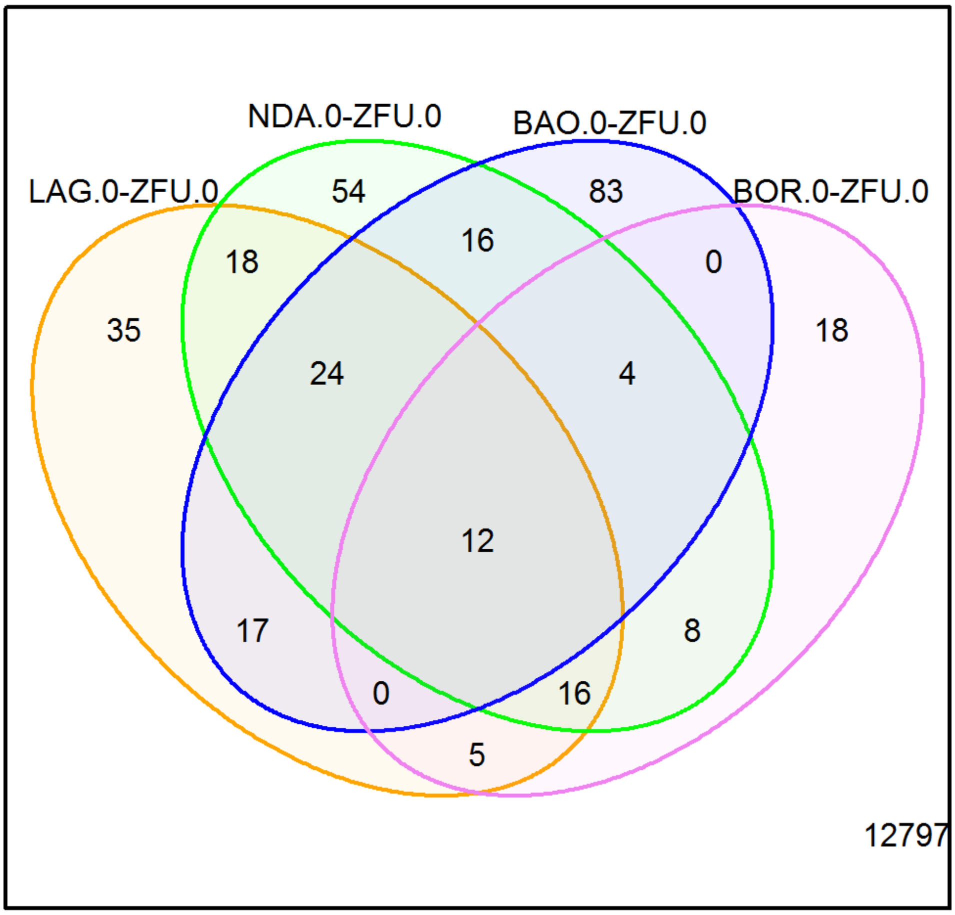


**S4 Figure. Venn diagram showing the intersection of genes identified as DE in the between-breed contrasts.**

Number of genes identified as DE in the four between-breed contrasts (LAG.0-ZFU.0, NDA.0-ZFU.0, BAO.0-ZFU.0, and BOR.0-ZFU.0) and at the intersections of the different sets.

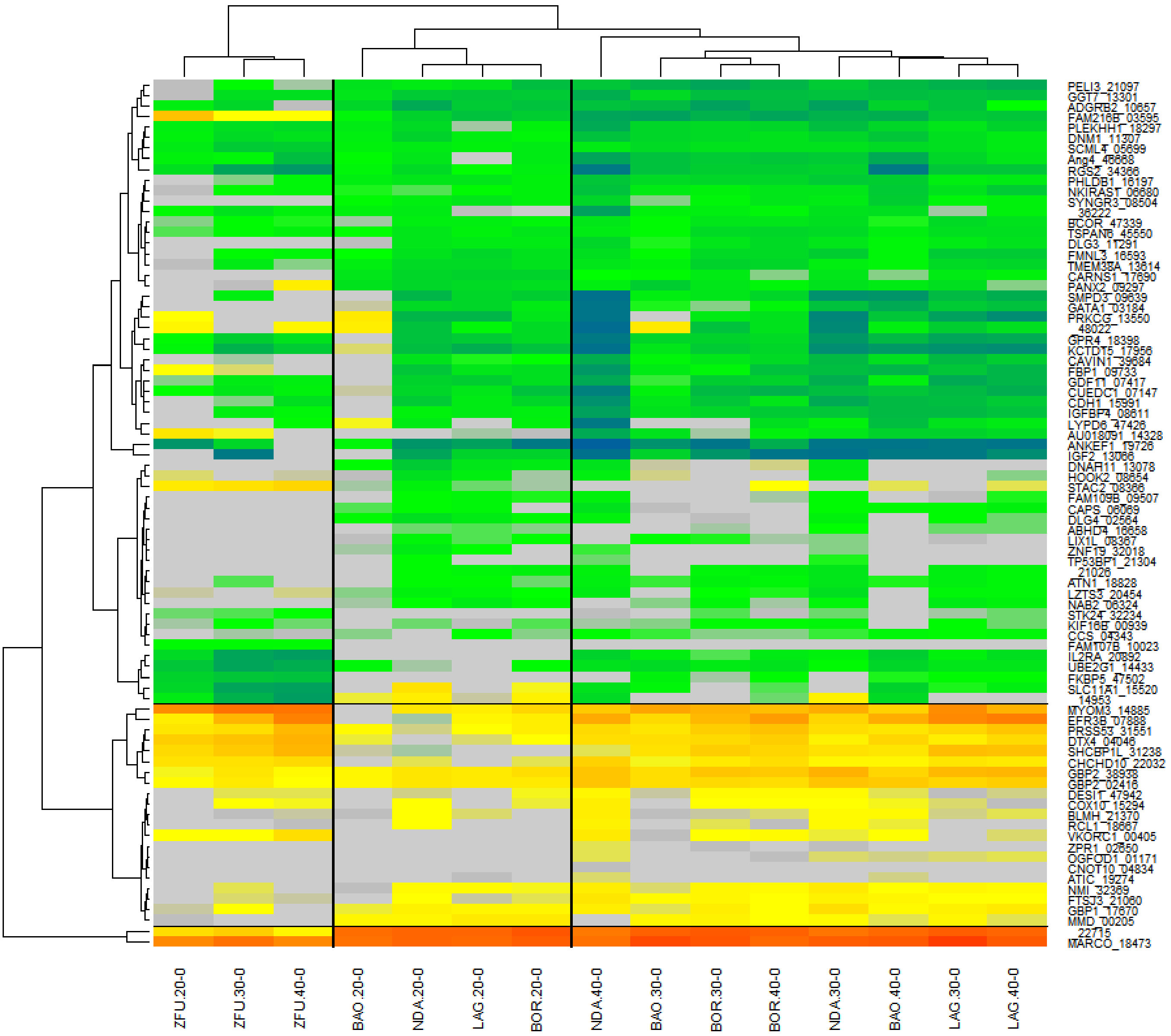


**S5 Figure. Heatmap on the logFC of a subset of 82 DEG.**

In columns are indicated the 15 within-breed contrasts ordered according to clustering, and in rows the genes identified as DE in the NDA.0-ZFU.0 contrast and either in ZFU contrasts (ZFU.20-0, ZFU.30-0, and ZFU.40-0) or in NDA contrasts (NDA.20-0, NDA.30-0, and NDA.40-0). Positive logFC, indicating up-regulated genes, are colored in warm colors from yellow to red, whereas negative logFC, indicating down-regulated genes, are colored in cold colors, from light green to blue. LogFC comprised between -0.20 and 0.20 are colored in grey.
